# Neutrophils regulate ITPR2 levels in epithelia by direct injection of elastase

**DOI:** 10.1101/2022.09.13.506588

**Authors:** N Ogino, M Fatima Leite, E Kruglov, H Asashima, DA Hafler, BE Ehrlich, MH Nathanson

## Abstract

The destructive role of neutrophils in inflammation is well known^1^ but they also have less damaging effects such as tissue remodeling and modulation of metabolism^2, 3^. Usually, neutrophils in tissues release toxic or digestive compounds into the extracellular region^4–8^. Here we report that neutrophils can inject their granule contents directly into hepatocytes. Neutrophil elastase within the hepatocytes selectively degrades the inositol trisphosphate receptor (ITPR), especially the type 2 isoform which is the predominant intracellular calcium release channel in these cells^9^. This action reduces calcium signals and cell proliferation without cellular damage. In response, the hepatocytes increase expression of serpins E2 and A3, which block the effect of elastase. This phenomenon is also observed in liver biopsies from patients with alcoholic hepatitis, a condition characterized by infiltration of neutrophils^10, 11^. This non-destructive and reversible effect on hepatocytes defines a previously unappreciated role of neutrophils in transiently regulating signaling mechanisms in epithelia.

---

ITPR2 accounts for 80% of the ITPRs pool in hepatocytes^9^ and is responsible for such varied calcium-mediated processes as secretion^12, 13^, gene transcription^14^ and cell proliferation^15–18^. ITPR2 is decreased in the liver in various inflammatory conditions^19^, which lead us to examine whether neutrophils directly affect ITPR2 levels and ITPR2-related calcium signaling. Co-culture with freshly isolated human neutrophils reduced ITPR2 in both primary human hepatocytes and the human liver-derived HepG2 cell line (Fig. 1A, B). Calcium signals were induced in HepG2 cells by stimulation with adenosine triphosphate (ATP), and the signals were markedly reduced in cells that were pre-incubated with neutrophils (Fig. 1C). Cell proliferation is calcium-mediated in the liver^15–18^, and this also was significantly attenuated in HepG2 cells that were co-cultured with neutrophils (Fig. 1D). The cytotoxic effect of neutrophils is well known^1, 5, 7^, but we found that most HepG2 cells co-cultured with neutrophils were alive rather than dead after 20 hours (Fig. 1E) and even after 40 hours (Fig. S1A). Furthermore, ITPR2 expression in the HepG2 cells was restored soon after removal of the neutrophils (Fig. 1F). Collectively, these findings show that neutrophils reversibly decrease ITPR2 and its associated calcium signaling in hepatocytes without causing cell death.

**Figure 1.**
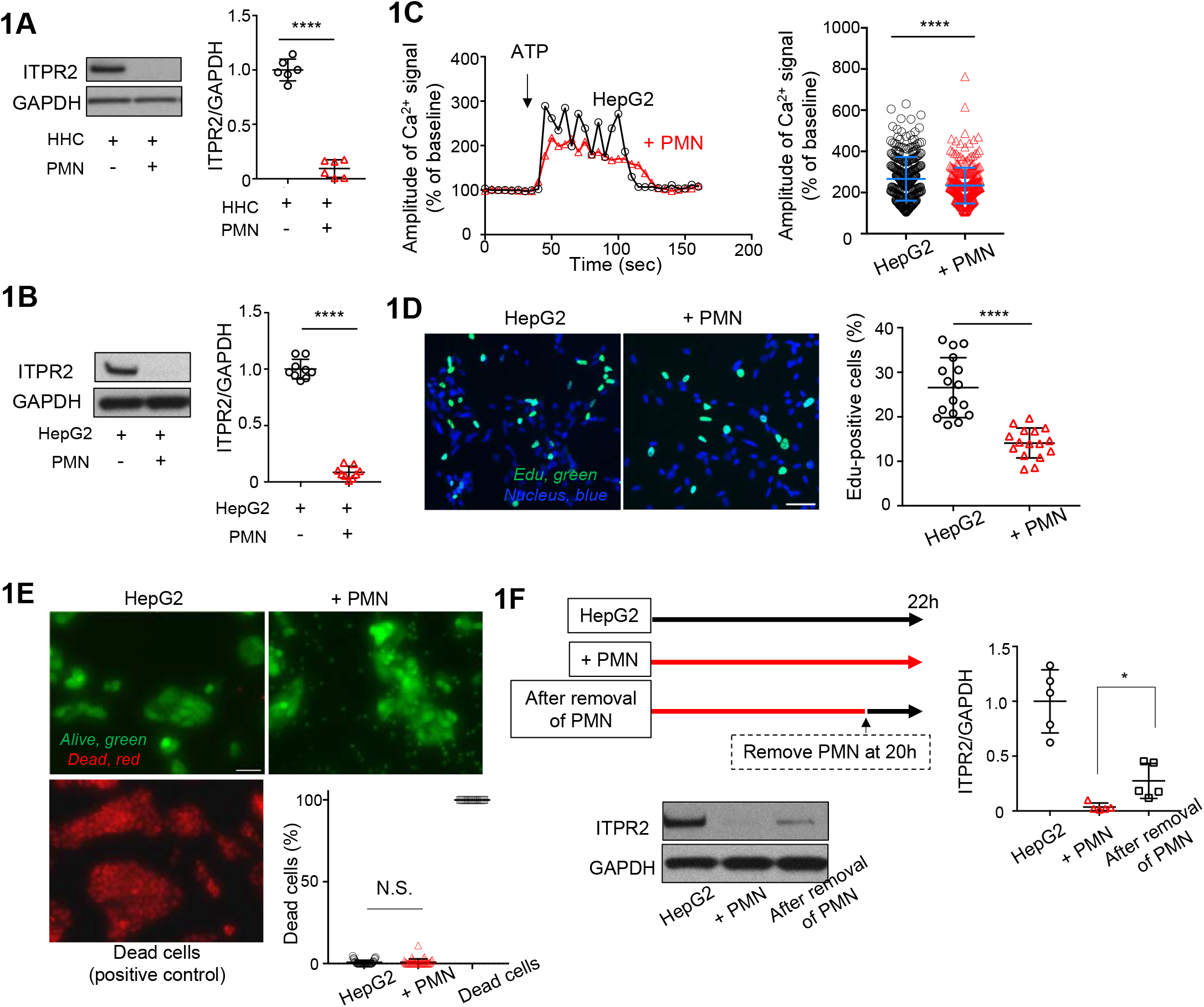
Neutrophils decrease ITPR2 levels, calcium signals and cell proliferation in hepatocytes but do not induce cell death. **(a)** Human polymorphonuclear neutrophils (PMN) decrease ITPR2 levels in primary human hepatocytes (HHC). HepG2 cells were co-cultured with PMN from 6 different healthy volunteers in a 1:1 ratio number for 18-24 hours. Following co-culture, hepatocytes were assessed for ITPR2 levels by immunoblotting. Representatives blots (*left*) and quantification (*right*) are shown. Data are mean ± SD, n=6 (****p<0.0001, unpaired t-test). **(b)** PMN decrease ITPR2 levels in HepG2 cells. Cells were co-cultured with PMN from 9 different volunteers in a 1:1 ratio number for 18-24 hours. Following co-culture, cells were assessed for ITPR2 levels by immunoblotting. Representatives blots (*left*) and quantification (*right*) are shown. Data are mean ± SD, n=9 (****p<0.0001, unpaired t-test). **(c)** PMN decrease the amplitude of Ca^2+^ signals evoked by adenosine triphosphate (ATP) in HepG2 cells. HepG2 cells pre-incubated for 20 hours with or without PMN were loaded with Fluo-4/AM (6 μM) for 30 minutes and then stimulated with ATP (20 μM). Left: Representative fluorescence intensity tracings from measurements in selected regions of interest of the cells. Right: Quantification from 9 coverslips of HepG2 cells (317 total cells) and 8 coverslips of HepG2+PMN (279 total cells), from 3 independent experiments for each condition. Graph shows mean ± SD (****p<0.0001, unpaired t-test). **(d)** PMN decrease proliferation of HepG2 cells. EdU (5-ethynyl-2’-deoxyuridine) proliferation assay was performed in HepG2 cells co-cultured with or without PMN in a 1:1 ratio number for 12 hours. Left: Representative image of cells with EdU-Alexa Fluor 488 labeling (green), indicative of proliferation, and Hoechst 33342 labeling of nuclei (blue). Scale bar, 50 μm. Right: Quantitative analysis of EdU-positive cells (%, n=5-6 coverslips for each condition in 3 independent experiments. Data are mean ± SD (****p < 0.0001, unpaired t-test). **(e)** PMN do not induce death of HepG2 cells. Live/dead cell analysis of HepG2 cells using double staining of calcein-AM (green) and Ethidium Homodimer-1 (EthD-1, red) with or without co-incubation with PMN for 20 hours. For positive control of dead cells, HepG2 cells were treated with 70% ethanol for 10 minutes. Representative image shows HepG2 cells stained with calcein-AM and EthD-1. Scale bar, 50 μm. Lower right panel shows the quantification of dead cells (%). Data are mean ± SD, n=5-10 fields in 3 cover slips per each condition in 3 independent experiments (N.S., not significant, unpaired t-test). **(f)** ITPR2 levels in HepG2 cells recover after PMN are removed. After co-culturing HepG2 cells with PMN for 20 hours, cells were washed with phosphate-buffered saline until PMN were no longer visible, then cultured for 2 hours and were collected (labeled as After removal of PMN). The ITPR2 levels was compared to that of HepG2 cells co-cultured with PMN for 22 hours or control (labeled as + PMN or HepG2 respectively). Representative blots (*left*) and quantification (*right*) are shown. Data are mean ± SD. n=3-5 (*p<0.05, unpaired t-test).

Several factors secreted by neutrophils are involved in inflammatory diseases of the liver and other organs and tissues^20, 21^. However, we found that ITPR2 was not decreased in HepG2 cells treated with neutrophil-conditioned medium (Fig. 2A) or in HepG2 cells separated from neutrophils by a semipermeable membrane (Fig. 2B). This unexpected preservation of ITPR2 suggests that neutrophils need to be in contact with HepG2 cells to affect ITPR2 levels. To confirm this, neutrophils were treated with phosphatidylinositol-specific phospholipase C (PI-PLC, 0.5 units/mL) to remove glycosylphosphatidylinositol (GPI)-anchored surface membrane proteins^22, 23^ before HepG2 cells were co-cultured with these neutrophils. The effect of these neutrophils was significantly attenuated compared to untreated neutrophils (Fig. 2C), even though viability of these neutrophils is not reduced^22^. Neutrophils can bind to integrins on some epithelia to activate signaling pathways^24, 25^, but blocking antibodies directed against integrins identified by pathway analysis of RNA-seq from HepG2 cells co-cultured with neutrophils (S2A, Supplementary Table 1), did not affect the reduction in ITPR2 that was seen in HepG2 cells co-cultured with neutrophils (Fig. 2D). In addition, neutrophil extracellular traps (NETs) can form in liver^7, 8^ and other organs^26^ to locally affect parenchymal cells. Typical mediators released by neutrophils include high mobility group box 1 protein (HMGB1), neutrophil elastase, and myeloperoxidase (MPO) ^26, 27^. However, extracellular administration of HMGB1, neutrophil elastase, or MPO to HepG2 cells had no effect on ITPR2 (Fig. 2E). Collectively, these findings provide evidence that neutrophils must be intact and in direct contact with HepG2 cells to decrease ITPR2. To further delineate the nature of the interaction between neutrophils and HepG2 cells, crude fractions of the neutrophils^28^ were administered to the HepG2 cells (Fig. 2F, S3A). ITPR2 was decreased by incubation with the granule-containing fraction but not by the cytoplasmic or plasma membrane fraction (Fig. 2G), and the effect of the granule fraction and homogenate were eliminated by boiling (Fig. 2G). Also, this effect of the granule fraction was not due to contamination by cell membrane proteins or nuclei (Fig. S3B). Furthermore, the granule fraction decreased ITPR2 in HepG2 cells in a concentration dependent manner, similar to the concentration-dependent effect of intact neutrophils on HepG2 cells (Fig. 2H). These findings suggest that neutrophil granule contents are required and that neutrophil membrane proteins alone are insufficient to decrease ITPR2 in HepG2 cells.

**Figure 2.**
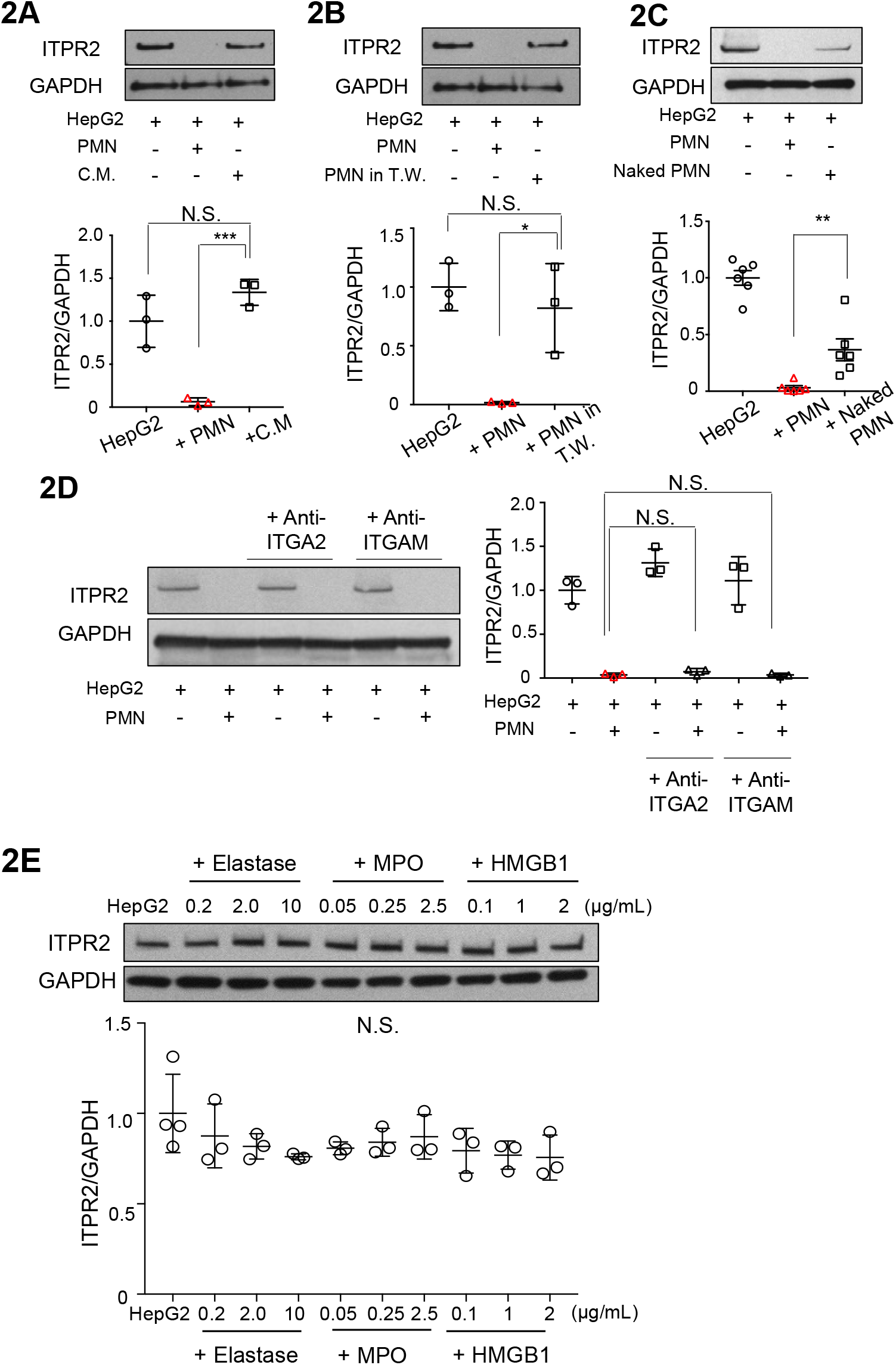
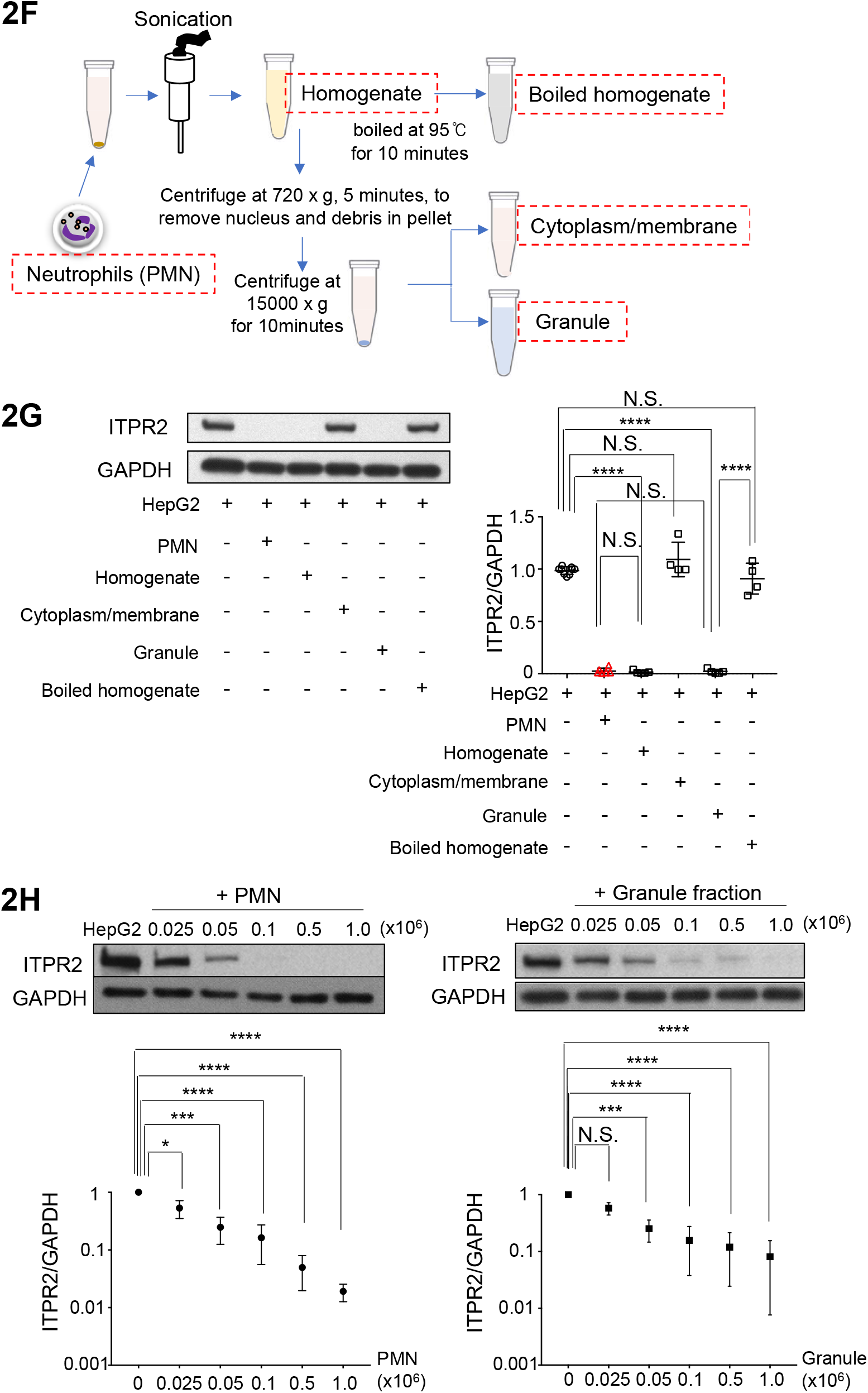
The direct contact and granule component of neutrophils contribute to the reduction of ITPR2 in HepG2 cells. **(a)** Conditioned medium (C.M.) from human polymorphonuclear neutrophils (PMN) do not decrease ITPR2 levels in HepG2 cells. Cells were cultured for 18-24h with C.M. derived from 16 hours incubation with PMN, and ITPR2 levels were assessed by immunoblotting. Representative blots (*top*), and quantification (*bottom*) are shown. Data are mean ± SD, n=3. (N.S., no significant difference, ***p<0.001 by one-way ANOVA). **(b)** PMN do not decrease ITPR2 levels in HepG2 cells that are separated by a semipermeable membrane. PMN were placed in the upper compartment of 3 μM pore transwell system (T.W.) and co-cultured with HepG2 cells cultured in the lower compartment. After 18–24 hours of co-culture, HepG2 cells were assessed for ITPR2 levels by immunoblotting. Representative blots (*top*) and quantification (*bottom*) are shown. Data are mean ± SD, n=3 (N.S., no significant difference, *p<0.05 by one-way ANOVA). **(c)** ‘Naked’ PMN partially lose the ability to decrease ITPR2 in HepG2 cells. Naked PMN were prepared by removing cell surface glycosylphosphatidylinositol (GPI)-anchored proteins by treatment with phosphatidylinositol-specific phospholipase C (PI-PLC). After co-culturing naked PMN with HepG2 for 18-24 hours, the ITPR2 levels in HepG2 cells was assessed by immunoblotting. Representative blots (*top*) and quantification (*bottom*) are shown. Data are mean ± SD, n=6 (**p<0.01, one-way ANOVA). **(d)** Functional blocking treatment by anti-integrin alpha-2 (ITGA2) or anti-integrin alpha M (ITGAM) antibodies do not prevent the decrease in ITPR2 of HepG2 cells by PMN. HepG2 cells were pre-incubated with anti-ITGA2 or ITGAM antibodies for 2 hours and then co-incubated with PMN. These are candidate integrins identified by bioinformatic analysis of RNA-seq from HepG2 cells after co-culturing with PMN. After co-culture, HepG2 cells were assessed for ITPR2 levels by immunoblotting. Representative blots (*left*) and quantification (*right*) are shown. Data are mean ± SD, n=3 (N.S., no significant difference by one-way ANOVA). **(e)** The components released in neutrophil extracellular traps (neutrophil elastase, myeloperoxidase (MPO) or high mobility group box-1 (HMGB1)) do not decrease ITPR2 levels in HepG2 cells. Cells were co-incubated with varying concentrations of elastase, MPO or HMGB1 for 20 hours and then assessed for ITPR2 levels by immunoblotting. Representative blots (*top*) and quantification (*bottom*) are shown. Data are mean ± SD, n=3-4 (N.S., no significant difference by one-way ANOVA). **(f)** Flowchart illustrating the experimental procedure to fractionate PMN is shown. **(g)** The granule fraction but not the cytoplasm/membrane fraction from PMN decreases ITPR2 in HepG2 cells. After incubation for 20 hours with the indicated fractions, HepG2 cells were assessed for ITPR2 levels by immunoblotting. Representative blots (*left*), and quantification (*right*) are shown. Data are mean ± SD, n=4 (N.S., no significant difference; ****p<0.0001 by one-way ANOVA). **(h)** The granule fraction decreases ITPR2 in HepG2 cells in a concentration-dependent fashion that follows first-order kinetics, similar to what is observed with intact PMN. HepG2 cells were co-cultured with PMN (*left*) or the granule fractions (*right*) extracted from the indicated number of PMN from 5 different healthy control subjects for 20 hours. Following co-culture, HepG2 cells were assessed for ITPR2 levels by immunoblotting. Representative blots (*top*) and quantification (*bottom*) are shown. Note both data sets are linear on a log-log scale. Shown are mean ± SEM, n=5 (N.S., not significantly different; *p<0.05; ***p<0.001; ****p<0.0001 by one-way ANOVA).

In order to identify the mediator of neutrophils’ effect, the time course was examined. ITPR2 levels in HepG2 cells were decreased when monitored at 1 hour after co-culture and persisted when measured 20 hours later (Fig. 3A). Calcium signaling in the HepG2 cells was also decreased after 1 hour of co-culture with neutrophils (Fig. S4A). Calcium signals showed significant recovery by 1 hour and further recovered by 19 hours after neutrophils were removed following 1 hour of co-culture (Fig. S4B). ITPR2 mRNA was increased rather than decreased in HepG2 cells 1 hour after co-culture (Fig. 3B), and ITPR2 immunoblots (Fig. 1B) subjected to longer exposure revealed a ladder-like appearance (Fig. S5A). These data suggest that the early decrease in ITPR2 is related to proteolysis rather than transcriptional regulation.

**Figure 3.**
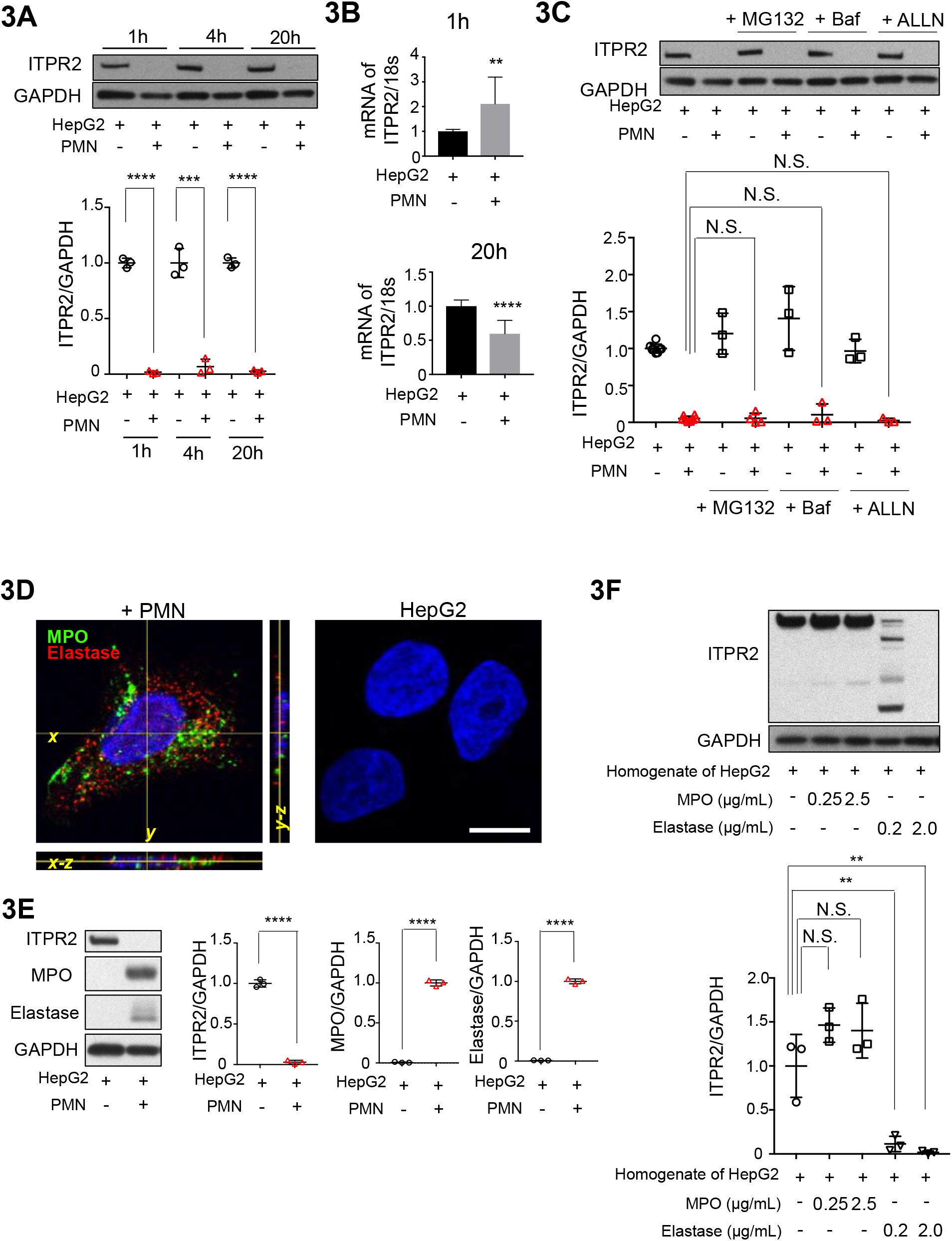
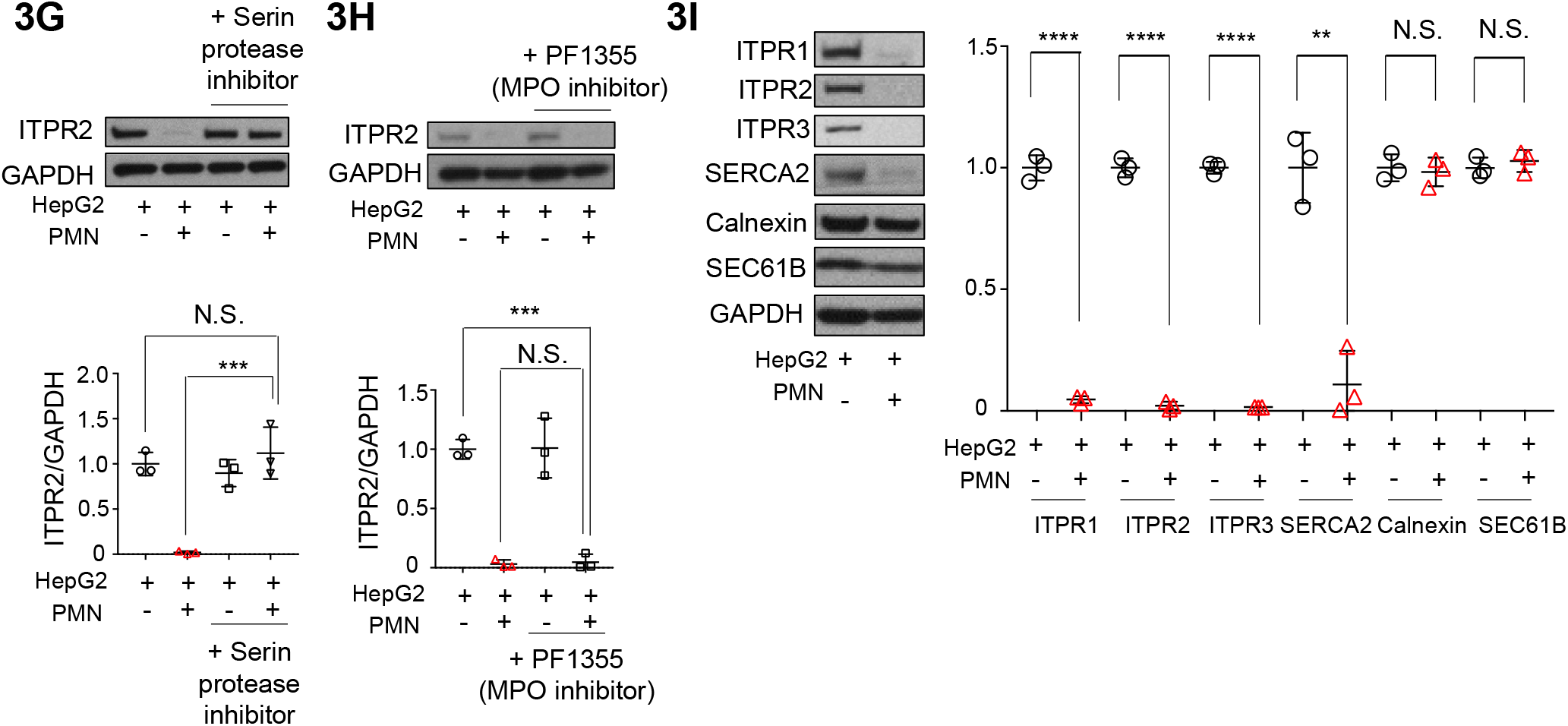
Neutrophils transfer elastase to HepG2 cells to degrade ITPR2. **(a)** The levels of ITPR2 in HepG2 cells are decreased 1 hour after administration of human polymorphonuclear neutrophils (PMN) and remains low after 4 and 20 hours. After co-culturing PMN with HepG2 cells for the indicated times, the amount of ITPR2 in HepG2 cells was assessed by immunoblotting. Representative blots (*top*) and quantification (*bottom*) are shown. Data are mean ± SD, n=3 (***p<0.001, ****p<0.0001, by one-way ANOVA). **(b)** ITPR2 mRNA in HepG2 cells is increased 1 hour after co-culture with PMN. Reverse transcription quantitative PCR (RT-qPCR) was performed to assess ITPR2 mRNA levels in HepG2 cells after co-culture with PMN for the indicated times. Quantitative analysis is shown. Data are mean ± SD, n=3 (**p<0.01, ****p<0.0001 by unpaired t-test). **(c)** The neutrophil-induced decrease in ITPR2 in HepG2 cells is not due to classical proteolytic mechanisms. HepG2 cells were pre-incubated with MG132 (proteasome inhibitor, 50 μM), bafilomycin A1 (Baf, autophagy inhibitor, 50nM), or ALLN (Ac-Leu-Leu-Nle-Aldehyde, calpain inhibitor, 50 μM) for 1 hour and followed for 1 hour after co-culture with PMN. The ITPR2 levels in HepG2 cells were assessed by immunoblotting. Representative blots (*top*) and quantification (*bottom*) are shown. Data are mean ± SD, n=3-4 (N.S., no significant difference by one-way ANOVA). **(d)** PMN transfer myeloperoxidase (MPO, *green*) and elastase (*red*) to HepG2 cells. Cells were co-cultured with PMN for 1 hour, washed with phosphate-buffered saline until PMN were not visible, then stained with anti-MPO and anti-elastase antibodies. Representative serial optical sections obtained by confocal immunofluorescence imaging (Z-stack) of HepG2 cells is shown. Hoechst 33342 is used to stain nuclei (blue). Scale bar, 20 μm. **(e)** Immunoblotting confirms that MPO and elastase are detected in HepG2 cells after co-incubated with PMN. After HepG2 cells were co-cultured with PMN for 1 hour, cells were washed with phosphate-buffered saline until no PMN were visible. Then, HepG2 cells were assessed for ITPR2, MPO and elastase by immunoblotting. Representative blots (*left*) and quantification (*right*) are shown. Data are mean ± SD, n=3 (****p<0.0001, unpaired t-test). **(f)** ITPR2 is degraded by elastase but not by MPO. HepG2 cells (3.0 x 10^6^) were homogenized in 100 μL of phosphate-buffered saline by sonication, followed by addition of MPO or neutrophil elastase at the concentrations indicated. Five minutes later, ITPR2 levels were assessed by immunoblotting. Note that there are multiple lower molecular weight bands in homogenate treated with the lower elastase concentration, but digestion is complete in homogenates treated with the higher concentration. Representative blot (*left*) and quantification (*right*) are shown. Data are mean ± SD, n=3 (N.S., no significant difference, **p<0.01 by one-way ANOVA). **(g)** The serin protease inhibitor AEBSF prevents the loss of ITPR2 in HepG2 cells induced by PMN. After co-culturing PMN with HepG2 cells with or without 10 μM AEBSF (4-(2-aminoethyl) benzenesulfonyl fluoride hydrochloride) for 1 hour, the ITPR2 levels in HepG2 cells was assessed by immunoblotting. Representative blots (*top*) and quantification (*bottom*) are shown. Data are mean ± SD, n=3. (N.S., no significant difference, ***p<0.001 by one-way ANOVA). **(h)** The MPO inhibitor PF1355 does not prevent the PMN-induced loss of ITPR2 in HepG2 cells. After co-culturing PMN with HepG2 cells with or without 10 μM PF-1355 (2-(6-(2,5-dimethoxyphenyl)-4-oxo-2-thioxo-3,4-dihydropyrimidin-1(2H)-yl) acetamide) for 1 hour, the ITPR2 levels in HepG2 cells was assessed by immunoblotting. Representative blots (*top*) and quantification (*bottom*) are shown. Data are mean ± SD, n=3. (N.S., no significant difference, ***p<0.001 by one-way ANOVA). **(i)** The target proteins in hepatocytes by neutrophil elastase are not non-specific. After co-culturing PMN with HepG2 cells for 1 hour, the amounts of ITPR1, ITPR2, ITPR3, SERCA2, calnexin, SEC61B in the HepG2 cells were assessed by immunoblotting. The ITPRs and SERCA2 were degraded but calnexin and SEC61B were not. Representative blots (*left*) and quantitative analysis (*right*) are shown. ITPR1, ITPR2 and ITPR3 have similar molecular weights, so these samples were blotted onto different membranes; GAPDH is checked at the same time. Data are mean ± SD, n=3. (N.S., no significant difference; **p<0.01; ****p<0.0001 by unpaired t-test).

Inhibitors of autophagy, proteasomes, or calpain did not prevent the decrease in ITPR2 (Fig. 3C). In addition, inhibitors of trypsin, and caspase-3, both of which can degrade ITPRs^29, 30^ did not prevent the decrease in ITPR2 (Fig. S5B). Therefore, we investigated whether specific neutrophil granule contents degrade ITPR2. After co-culturing HepG2 cells with neutrophils for 1 hour and washing to ensure that no neutrophils remained, confocal immunofluorescence staining revealed that MPO and neutrophil elastase were contained within the HepG2 cells (Fig. 3D). This was confirmed by Western blot (Fig. 3E). The presence of MPO was also confirmed by Western blot in human hepatocytes after co-culture with neutrophils (Fig. S5C). Moreover, when HepG2 cells were incubated with the red-stained granule fraction, confocal microscopy revealed that granules were incorporated into the cells (Fig. S5D) ^31^. This was confirmed by immunofluorescence staining with MPO antibody (Fig. S5D). Next, HepG2 cell homogenates were treated with MPO or elastase, and elastase but not MPO degraded ITPR2 (Fig. 3F). Moreover, in HepG2 cells co-cultured with neutrophils, the decrease in ITPR2 was completely blocked by 4-(2-aminoethyl) benzenesulfonyl fluoride hydrochloride (AEBSF), a serine protease inhibitor, but not by either of two MPO inhibitors (Fig. 3G, 3H, S5E). To understand the specificity of elastase in this setting, we examined several ER proteins and found that the other two ITPR isoforms and SERCA2 also were targets. However, SEC61B and calnexin were not (Fig. 3I). These findings collectively suggest that neutrophils transfer elastase into HepG2 cells to selectively but transiently degrade ITPR2 and several other proteins, with a focus on proteins relevant for calcium signaling.

The observations that neutrophils are not damaging HepG2 cells in this system (Fig. 1E, S1A) and that cells recover when neutrophils are removed (Fig. 1F, S4B) suggest that hepatocytes have a mechanism to temporally limit this effect of neutrophils. To investigate this pathway for preservation, bulk RNA sequencing was performed on HepG2 cells 20 hours after incubation with either neutrophils or their granule fraction (Fig. 4A). We identified 101 genes upregulated and 108 genes downregulated by neutrophils, and 50 genes upregulated, and 49 genes downregulated by the granule fraction (Fig. 4B). Of these groups of genes, 10 were altered by both neutrophils and their granules, including serpin E2 and serpin A3 (Fig. 4B, S6). Therefore, we investigated whether the proteins encoded by these two genes may suppress the effect of neutrophil elastase on ITPR2. RT-qPCR of mRNA in HepG2 cells 1 h after neutrophil treatment showed that both of serpin E2 and serpin A3 were significantly elevated (Fig. 4C). In addition, administration of recombinant serpin E2 or serpin A3 inhibited the degradation of ITPR2 by elastase in HepG2 cell homogenates (Fig. 4D). These findings provide evidence that hepatocytes increase expression of serpin E2 and serpin A3 in order to limit the duration of the effects of the elastase that is inserted into the cells by neutrophils.

**Figure 4.**
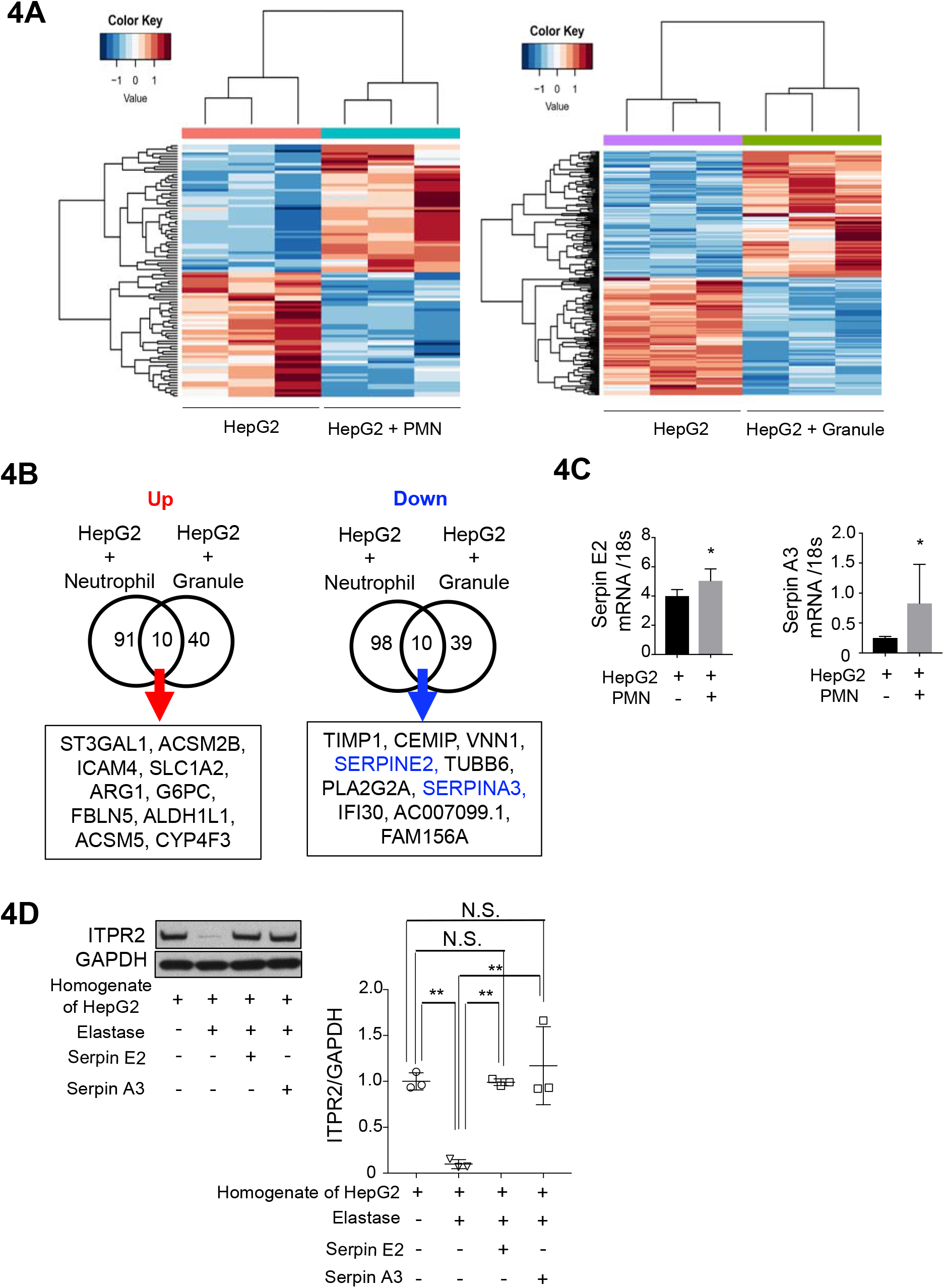
Hepatocytes respond to neutrophil elastase by altering the expression of serpin E2 and serpin A3. **(a)** Heat maps are shown for two experiments, comparing RNA-seq data of HepG2 cells treated with human polymorphonuclear neutrophils (PMN) or extracted granule fraction (both from three different volunteers) for 20 hours and respective controls (HepG2). The left map shows the PMN treatment, and the right one shows the granule fraction treatment. **(b)** The Venn diagram shows the number of genes that were significantly up- or down-regulated (false discovery rate (FDR) < 0.05, log fold change < −0.5 or >0.5) for HepG2 cells treated with neutrophil or granule fraction compared to the respective controls. A list of genes commonly up- or down-regulated in the two RNA-seqs is shown in the bottom. 10 genes were extracted, including Serpin E2 and Serpin A3. **(c)** The mRNA levels of serpin E2 and serpin A3 in HepG2 cells are significantly elevated after co-culture with PMN for 1 hour. Reverse transcription quantitative PCR (RT-qPCR) was performed to assess the mRNA levels of serpin E2 and serpin A3 in HepG2 cells after co-culture with PMN for 1 hour. Data are mean ± SD, n=3. (*p<0.05, unpaired t-test). **(d)** Degradation of ITPR2 by neutrophil elastase is inhibited by serpin E2 and serpin A3. HepG2 cells (1.0 x 10^6^) were homogenized by sonication in 100 μL phosphate-buffered saline followed by addition of neutrophil elastase (0.02 μg/mL) with or without serpin E2 or serpin A3 (20 μg/mL). 5 minutes later, ITPR2 levels were assessed by immunoblotting. Representative blots (*left*) and quantification (*right*) are shown. Data are mean ± SD, n=3. (N.S., no significant difference, **p<0.01 by one-way ANOVA).

Finally, we investigated whether the effect of neutrophils on epithelia was liver-specific, and whether it occurred in human disease. Parenchymal infiltration of neutrophils can occur in most organs^3, 24^, so epithelial cell lines derived from colon, pancreas, and lung were co-cultured with neutrophils for 1 hour to measure levels of granule proteins and ITPR2. Both MPO and elastase were detected and ITPR2 was decreased in HCT166, A549 and PANC-1 cells (Fig. 5A). To determine the relevance of our observations for human disease, we compared liver biopsies from patient specimens that were histologically normal to specimens from patients with biopsy-proven alcoholic hepatitis, which is characterized in part by infiltration of neutrophils^5, 11^. Immunochemical staining for ITPR2 was significantly decreased in hepatocytes in the biopsies from patients with alcoholic hepatitis, relative to histologically normal controls (Fig. 5B). Similarly, ITPR2 as detected by immunoblot was significantly decreased relative to controls in liver homogenates from mice in which alcoholic hepatitis had been induced using the NIAAA mouse model (Fig. S7) ^32^. Finally, immunohistochemical staining for neutrophil elastase was performed on human liver biopsy specimens. In the normal liver, this staining identified occasional neutrophils, several which appeared to be present in the sinusoids. In alcoholic hepatitis biopsies, on the other hand, significantly more elastase-positive cells were found, many of which were infiltrating the liver parenchyma (Fig. 5C). Moreover, higher power images identified elastase-positive granules in hepatocytes that were in proximity to neutrophils (Fig. 5D). Collectively, these findings provide evidence that neutrophils can insert elastase into epithelia in a variety of organs, and that this occurs in human disease.

**Figure 5.**
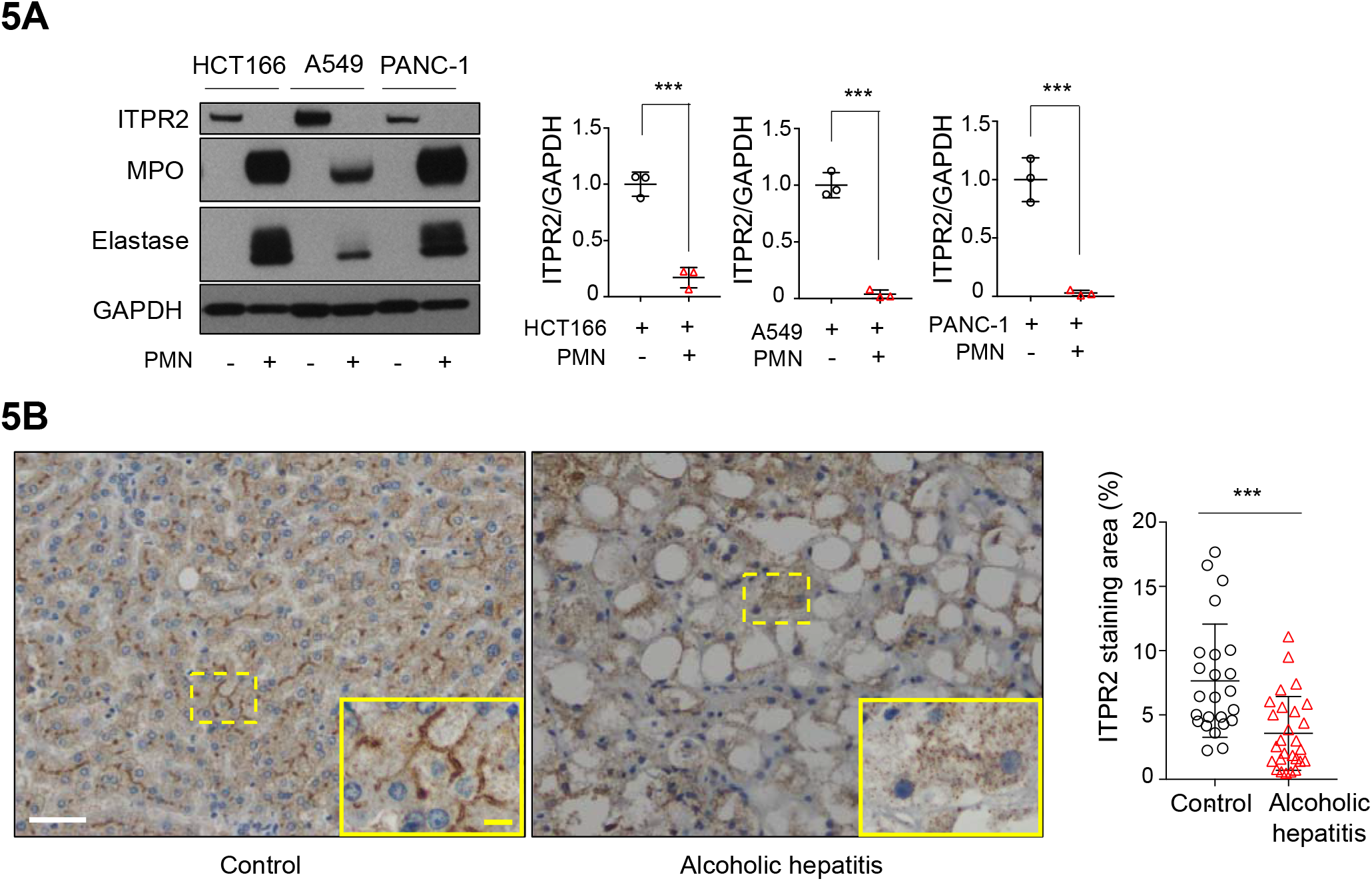
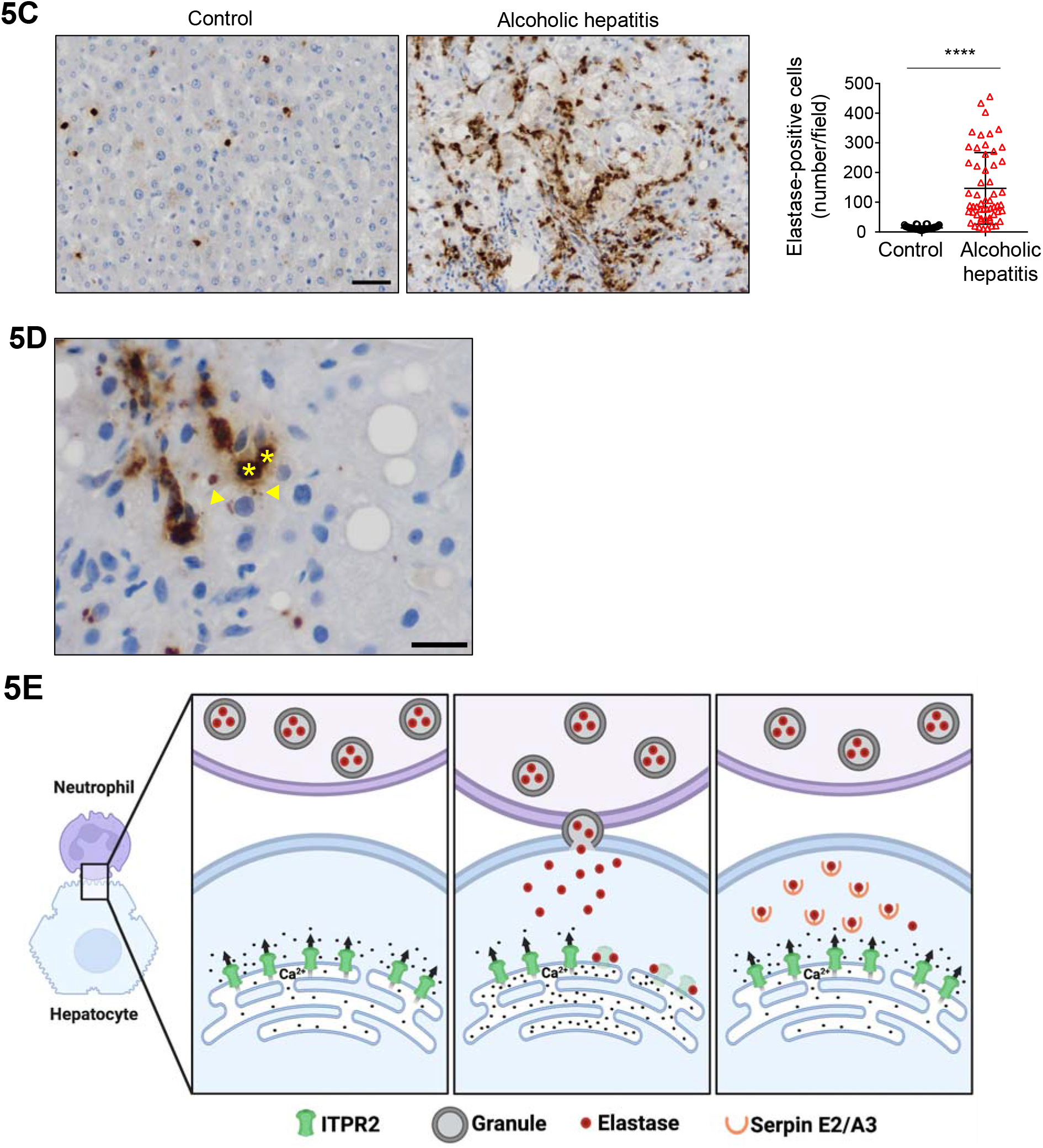
Degradation of ITPR2 by tissue-infiltrating neutrophils occurs in patients with alcoholic hepatitis but is not liver-specific and occurs in other epithelia. **(a)** Neutrophils transfer elastase and eliminate ITPR2 in cells derived from colon, lung, and pancreas. Human polymorphonuclear neutrophils (PMN) were co-cultured with HCT166 (colon), A549 (lung), and PANC-1 (pancreas) cells for 1 hour. After co-culture, ITPR2, myeloperoxidase (MPO) and elastase in these cells were assessed by immunoblotting. Representative blots (*left*) and quantification for ITPR2 (*right*) are shown. Data are mean ± SD, n=3 (***p<0.001, unpaired t-test). **(b)** ITPR2 staining in hepatocytes is reduced in liver biopsy specimens from patients with alcoholic hepatitis (AH) relative to histologically normal controls. Representative immunochemistry images (*left*) and quantitative analysis of ITPR2 staining (*right*) are shown. The brown-colored area is stained with anti-ITPR2 antibody. The yellow areas are under high magnification. Scale bar, 50 μm (white), 10 μm (yellow). Data are mean ± SD; 3 fields were quantified in each biopsy specimen, which included 8 controls and 9 from patients with AH (***p<0.0001 using one-tailed unpaired t-test). **(c)** Elastase-positive neutrophils are much more prevalent in liver biopsy tissue from AH patients than in normal controls. Control and AH liver tissue were immunohistochemically stained with anti-neutrophil elastase antibodies. Scale bar, 50 μm. The number of elastase-positive cells was markedly increased in liver tissue from AH patients. Representative immunohistochemical staining (*left*) and the number of elastase-positive cells in each field using 20 x magnification (*right*) is shown. Cells were counted in 5-7 areas in each specimen (n=8 control and n=9 AH specimens; ****p<0.0001 by unpaired t-test). **(d)** Neutrophil elastase is present in hepatocytes that are in proximity to neutrophils. Shown is a representative image of immunochemical staining for elastase in a liver biopsy from a patient with AH. *Asterisks* indicate neutrophils and *arrowheads* denote elastase staining in hepatocytes (60x magnification, scale bar, 20 μm). **(e)** Sequence of events in neutrophil-hepatocyte interactions. *Left:* neutrophils are in proximity to hepatocytes; *Center:* neutrophils directly contact hepatocytes to transfer granule contents into the hepatocytes. The neutrophil elastase degrades ITPR2; *Right:* The neutrophil moves away from the hepatocyte, which recovers by producing serpin E2 and A3 to block the elastase.

Calcium signaling regulates a wide range of cellular function, and in epithelia this is mediated mostly by calcium release from ITPRs^33^. ITPR2 is the predominant isoform in hepatocytes^9^, where it plays an important role in secretion by promoting targeting and apical insertion of transporters for bile secretion^12, 22^ and in liver regeneration by promoting cell proliferation^15–18^. Consequently, liver diseases in which ITPR2 is reduced result in impairments in bile secretion^12^ and liver regeneration^19^. Both bile secretion and liver regeneration are impaired in alcoholic hepatitis as well^34, 35^, consistent with our observation that ITPR2 also is reduced in this clinical condition (Fig. 5B). Cells employ a range of strategies to regulate their own levels of ITPRs in order to control their calcium signals, including transcriptional^36^ and epigenetic^14^ mechanisms, plus a variety of degradative mechanisms^37^. ITPR levels also can be regulated by neighboring cells. For example, ITPR3 is the dominant isoform in cholangiocytes^38^, which is another epithelial cell type in liver, and its expression in these cells can be reduced by circulating endotoxin that binds to TLR4 on their cell membrane^39^ or by neutrophils, which bind to integrins on their membrane^22^. The current work provides a different mechanism for neighboring cells to modulate ITPRs in target cells, by direct insertion of a degradative enzyme (Fig. 5E). It appears to be particularly important that this neutrophil-hepatocyte interaction does not reduce viability of the hepatocyte (Fig. 1E, S1A), that the effects of neutrophil elastase have some specificity (Fig. 3I), and that the hepatocyte responds by expressing serpins to eliminate the elastase at the end of the interaction (Fig. 4C). These three features together allow the neutrophil to transiently but specifically modulate hepatocyte behavior in response to hepatic inflammation. This interaction may in fact be beneficial, because increased infiltration of neutrophils into the liver is correlated with increased rather than decreased survival in patients with alcoholic hepatitis^40^, an observation that was previously puzzling. These insights define new ways in which neutrophils can interact with parenchymal cells in a non-destructive fashion, as well as new ways in which epithelia can modulate their calcium signals in space and time to adapt to changes in their tissue environment.

## METHODS

### Ethics statement

Liver biopsy specimens from patients with alcoholic hepatitis (based on clinical presentation plus histological diagnosis) or histologically normal control specimens from surgical resections, plus blood samples from healthy volunteers were obtained at Yale-New Haven Hospital (YNHH) in New Haven, CT. This study was conducted under the auspices of protocols approved by the Institutional Review Board on the Protection of the Rights of Human Subjects (Yale University). The Human Investigation Committee protocol numbers are HIC-2000025846 and HIC-1304011763. Written informed consent was obtained from all participants. The study protocol conformed to the ethical guidelines of the Helsinki Declaration.

### Isolation of neutrophils from human blood samples

Peripheral venous blood samples were collected in K2 EDTA tubes (Becton, Dickinson and Company, Franklin Lakes, New Jersey, USA, 366643) from healthy human volunteers not taking drugs or alcohol. Human neutrophils were isolated and purified as previously described^22^. Using density gradient centrifugation with PolymorphPrep (Proteogenix, Schiltigheim, France, AN1114683) according to the manufacturer’s instructions, blood was layered in PolymorphPrep solution and centrifuged at 20 °C, 500 x g for 35 minutes without brakes. After centrifugation, the neutrophil layer was collected and washed with diluted N-2-hydroxyethylpiperazine-N’ 2-ethanesulfonic acid (HEPES) buffered saline to remove PolymorphPrep residue. Neutrophils were collected by centrifugation at 400 x g for 10 minutes. To purify neutrophils and remove residual erythrocyte contamination, cells were washed with ACK lysis buffer (Lonza, Bend, Oregon, USA, BP10-548E). Neutrophils were harvested by centrifugation and resuspended in the medium used for the cells to be administered. Cell density was quantified using a hemocytometer. Purity of neutrophil preparations was confirmed by Wright staining or May-Grunwald-Gemza staining/Pappenheim staining. Only isolates with >95% neutrophils were used. To remove GPI-anchored membrane protein, neutrophils were treated with phosphatidylinositol-specific phospholipase C (PI-PLC, 0.5 units/mL, Sigma-Aldrich, P5542) at 37°C for 30 minutes with gentle stirring as previously described^22^.

### Human liver specimens and histological examination

Alcoholic hepatitis specimens were from patients treated at YNHH from 2012 to 2014 with a clinical and histological diagnosis of alcoholic hepatitis; clinical characteristics of patients with biopsy-proven alcoholic hepatitis at YNHH were analyzed. Histologically normal tissue was obtained from the peripheral portion of liver resections performed for metastatic colorectal cancer, and cases were identified by review of pathology reports at YNHH from 2010 to 2017.

### Immunohistochemistry staining for ITPR2 and elastase

Immunohistochemical staining with anti-ITPR2 antibody (a gift from Dr. Richard Wojcikiewicz, SUNY, Syracuse, NY, USA) and analysis of ITPR2 was performed in liver biopsies from histologically normal livers and from patients with alcoholic hepatitis as previously described^19^. Staining by anti-neutrophil elastase (Abcam, ab131260) was performed in the same way using Novo Link Polymer Detection System (Leica biosystems, RE7290-K).

### Calculation of the number of elastase-positive cells

To identify and quantify the number of immune cells positive for neutrophil elastase staining, images were taken from 4-5 different regions of each liver biopsy or liver tissue specimen using a digitized Olympus BX51 microscope with a 20x objective. Representative images of elastase-positive granules in hepatocytes were taken at 60x.

### Cell culture and reagents

Human hepatocytes were obtained from the Liver Tissue Procurement and Distribution System of the National Institutes of Health (University of Pittsburgh), and maintained on collagen-coated plates using Hepatocyte Basal Medium (Lonza, CC-3197) with dexamethasone and insulin (each 0.1 μM, Millipore-Sigma, D4902 and I2643 respectively) and overlaid with Matrigel (Corning, 356237) as previously described^41^. HepG2 cells were obtained from American Type Culture Collection (ATCC, Manassas, VA, USA). HepG2 cells were cultured in Eagle’s Minimum Essential Medium (EMEM, ATCC, 30-2003) supplemented with 10% heat-inactivated fetal bovine serum (Gibco, 10438026), 100 U/ml penicillin G and 100 μg/ml streptomycin (Gibco, 15140122), at 37°C, 5% CO2. HCT166, A549 and PANC-1 cells were obtained from Dr. Ikki Sakuma (Chiba Univ.), from Dr. Samir Gautam (Yale Univ.) and from Dr. Moitrayee Bhattacharyya (Yale Univ.), respectively, which had been purchased from ATCC. These cells were maintained in culture in Dulbecco’s Modified Eagle’s Minimum Essential Medium (DMEM, ATCC, 30-2002) supplemented with 10% heat-inactivated fetal bovine serum and 100 U/ml penicillin G, and 100 μg/ml streptomycin, at 37 °C, 5% CO2. For experiments using adherent cells, one million cells were plated and used for experiments the following day, unless otherwise noted. For experiments using human neutrophils, the ratio of neutrophils to adherent cells was 1:1 unless otherwise noted.

HepG2 cells were treated with neutrophil elastase (Millipore-Sigma, 324681), neutrophil myeloperoxidase (MPO, Millipore-Sigma, 475911), or high mobility group box-1 (HMGB1, R&D systems, 1690-HMB) at the indicated concentrations for 20 hours. MG132 (Millipore-Sigma, 474790) was used at 50 μM to inhibit proteasomes. Bafilomycin A1 (Sigma Aldrich,19-148) was used at 50 nM to inhibit autophagy. N-Acetyl-L-leucyl-L-leucyl-L-norleucinal (ALLN, Millipore-Sigma, 208719) was used at 50 μM to inhibit calpain. These were administrated to HepG2 cells as pretreatment for 1 hour. A cell permeable caspase3 inhibitor (Millipore-Sigma, 235423) was used at 50 μM for 20 hours. AdaAhx3L3VS (Adamantane-acetyl-(6-aminohexanoyl)3-(leucinyl)3-vinyl-(methyl)-sulfone), a cell permeable trypsin inhibitor (Millipore-Sigma, 114802) was used at 50 nM for 20 hours. AEBSF, a cell permeable serin protease inhibitor (4-(2-Aminoethyl) Benzenesulfonyl Fluoride Hydrochloride, MP Biochemicals, 02193503-CF) was used at 10 μM. MPO inhibitor 2 (4-(5-Fluoro-1H-indol-3-yl) butanamide, Millipore-Sigma, 5049080001) was used at 5 μM and PF1355 (Cayman Chemicals, 22222) was used at 10 μM.

### Neutrophil fractionation

The homogenate fraction was obtained by centrifuging neutrophils in medium at 400 x g 5 minutes and treating the pellet with a stick ultrasound device (Fisherbrand™ CL18) at 120 watts, 20 kHz output for 3 seconds, three times. This was subjected to centrifugation at 720 x g 5 minutes and the settled pellet (debris and nuclei) was discarded. The supernatant was centrifuged at 15,000 x g for 10 minutes. The pellet was used as the granule fraction and the supernatant as the cytoplasmic or membrane fractions^28^. To extract purer plasma membrane protein components, cell plasma membrane and nuclei were extracted from neutrophils using the Minute™ Plasma Membrane Protein Isolation and Cell Fractionation Kit (Invent Biotechnologies Inc, Plymouth, MN, SM-005) according to the manufacturer’s instructions.

### Cell proliferation assay

For image-based detection of proliferating cells, HepG2 cells were plated onto coverslips in 6-well plates, then incubated with neutrophils for 12 hours. Fluorescent labeling of proliferating cells was detected using the Click-iT EdU Alexa Fluor 488 Imaging kit (Invitrogen, C10637) according to the manufacturer’s instructions. Five to six pictures were taken using a Zeiss Axio Observer epifluorescence microscope with 20x objective lens per coverslip. Overall cell numbers were measured by nuclear staining with Hoechst 33342 and the percentage of cells that had taken up 5-ethynyl-2’-deoxyuridine (EdU) was assessed.

### Cell death confirmation assay

For image-based measurement of cell death, HepG2 cells were plated onto coverslips in 6-well plates. To determine whether neutrophils induce cell death of HepG2 cells, dead cells were detected using a LIVE/DEAD Viability/Cytotoxicity kit (Molecular Probes, L3224), which uses calcein-AM and ethidium homodimer-1 (EthD-1) as markers. HepG2 cells with or without neutrophil co-culture for 20 or 40 hours were double stained. For positive control for dead cells, HepG2 cells were treated with 70% ethanol for 10 minutes. Five to six images were obtained using a Zeiss Axio Observer epifluorescence microscope with 20x objective lens per coverslip. The overall cell count was determined by adding the live cells stained with calcein-AM to the dead cells stained with EthD-1, but also checking nuclear staining with Hoechst 33342 to assess the percentage of dead cells.

### Neutrophil-conditioned medium and neutrophil co-culture in a transwell system

Human neutrophils were cultured in EMEM medium for 16 hours at 37°C. Neutrophils were removed by centrifugation (400 x g, 3 minutes) and the supernatant was used as the neutrophil-conditioned medium. HepG2 cells were incubated with the neutrophil-conditioned medium for 18-24 hours. Cells were harvested and assessed for ITPR2 levels. Neutrophils were placed in the upper compartment of a 3 μM pore transwell system (FALCON-353096) and co-cultured with HepG2 cells in the lower compartment. After 18-24 hours of co-culture, HepG2 cells were harvested and assessed for ITPR2 levels by immunoblotting.

### Functional blocking interventions

To inhibit the function of Integrin alpha M (ITGAM) or integrin alpha-2 (ITGA2) in HepG2 cells, an anti-ITGAM antibody (Thermo Fisher Scientific, # 14-0112-82, clone M1/70) or anti-ITGA2 antibody (Santa Cruz Biotechnology, # sc-53502, clone P1E6) were used as previously described^22, 42, 43^. In brief, HepG2 cells were seeded overnight and preincubated with ITGAM antibody (5 μg/mL) or ITGA2 antibody (10 μg/mL) for 2 hours at 37°C. The antibodies were removed, and cells were rinsed with phosphate-buffered saline (PBS) and co-cultured with or without neutrophils for an additional 18 hours. Cells were then harvested and assessed by immunoblotting.

### Confocal Fluorescence Imaging

Confocal immunofluorescence was performed as previously described^19^. HepG2 cells were plated onto glass coverslips (Thermo Fisher Scientific, 12542B) and incubated with neutrophils for 1 hour. The cells were fixed with 4% paraformaldehyde (Electron Microscopy Science, 15710) and permeabilized with 0.5% Triton X-100 (Sigma-Aldrich, X100). After washing in PBS, non-specific binding was blocked using 1% bovine serum albumin (BSA, Sigma-Aldrich, A2153), 0.05% Tween in PBS for 30 minutes at room temperature. For co-immunolabelling with primary antibodies, rabbit anti-myeloperoxidase (MPO) antibody 1(Abcam, 1:100 dilution, ab9535) and mouse anti-neutrophil elastase antibody (R&D systems, 1:100 dilution, MAB91671) were incubated for 8 hours, at 4°C, followed by a 1 hour incubation at room temperature with goat anti-mouse Alexa 564 and goat anti-rabbit Alexa Fluor 488 secondary antibodies (1:500; Invitrogen, A-11030 and A-11008 respectively), respectively. DNA was stained with DAPI or Hoechst 33342 (1:1000, Invitrogen, D1306 and H3570 respectively). After labelling with fluorochromes, the glass coverslips were mounted in an antifade fluorescent mounting medium (Electron Microscopy Sciences, 17966). Images were then obtained using a ZEISS LSM 710 or 880 confocal fluorescence microscope (Carl Zeiss, New York, USA).

### Granule staining and cellular uptake

CellMask Orange fluorescent lipophilic membrane dye (Thermo Fisher Scientific, C10045) was used to study the uptake of granule fraction constituents in HepG2 cells as previously described^31^. First, the granule fraction from human neutrophil (3.0 x 10^6) were stained with 7.5 μg/ml CellMask dye diluted in PBS in a final volume of 200 μl. The samples were incubated for 10 minutes at 37°C. Three centrifugations of 10 minutes at 15,000 x g were performed with the addition of PBS for each to wash the sample thoroughly and remove unbound dye. Pellet was resuspended with EMEM medium and added for HepG2 cells. After incubation for 1 hour in a conventional tissue culture incubator (37°C, 5% CO2), a Hoechst 33342 was used to label the nuclei at 5 μg/ml. Five images of living cells in each group were then acquired on a LSM710 confocal microscope using 40× magnification objectives and 543 nm lasers. In addition, HepG2 cells seeded on coverslips were treated with unstained granule fraction for 1 hour, fixed with 4% paraformaldehyde, and stained with MPO antibody using the immunostaining method described in the previous section and observed.

### Calcium signaling

Cytosolic calcium signals were measured as previously described^44^. Fluorescence intensity of Fluo4/AM (Invitrogen, F14201) in HepG2 cells in response to adenosine triphosphate (ATP, Millipore-Sigma, 20-306) was monitored with a Zeiss LSM 710 confocal microscope. Briefly, cells attached to glass coverslips were incubated with HEPES buffer (pH 7.4), then loaded with the calcium-sensitive fluorescent dye Fluo-4/AM (6 μM) for 30 minutes at 37°C. Coverslips containing the labelled cells were transferred to a custom-built perfusion chamber on the stage of the confocal microscope, and the cells were perfused with the HEPES buffer while stimulated with 20 μM ATP, which is a potent agonist of G protein-coupled receptors in these cells. Relative Fluo-4 fluorescence intensity after stimulation was analyzed in selected regions of interest using ImageJ (National Institutes of Health) and compared with baseline Fluo-4 fluorescence intensity.

### RNA isolation and quantitative real-time PCR

Total RNA was extracted and purified by RNeasy Mini Kits (QIAGEN, 74104) according to the manufacturer’s protocol. One half microgram of total RNA was reverse transcribed into cDNA using an iScript cDNA Synthesis kit (Bio-Rad, 1708891). All TaqMan primers and probes were obtained from Thermo Fisher Scientific: *ITPR2* (Hs00181916_m1), *Serpin E2* (Hs00299953_m1), *Serpin A3* (Hs00153674_m1), 18S ribosomal RNA (rRNA, Hs03003631_g1). Real-time PCR reactions were performed in triplicate using an ABI 7500 Sequence Detection System (Applied Biosystems). The expression of target genes was normalized to 18S rRNA and quantification of relative expression was determined as previously described^22^.

### Immunoblotting analysis

Western blot analysis of lysates was performed as previously described^22^. The following primary antibodies were used: ITPR1 (Alomone labs, ACC-019), ITPR2 (Santa Cruz Biotechnology, 398434), ITPR3(BD Transduction Laboratories, 610313), SERCA2 (Cell signaling, 4388), Calnexin (Abcam, ab92573), SEC61B (Thermo Fisher Scientific, PA3-015), MPO (Fisher Scientific, RB373A0), elastase (R&D systems, MAB91671), and GAPDH (Thermo Fisher Scientific, Clone 6C5, AM4300). The band intensities of proteins of interest were quantified and analyzed using ImageJ software.

### Bulk RNA-seq and Ingenuity Pathway Analysis

cDNA and library preparation and sequencing: RNA was isolated using RNeasy Mini Kits (QIAGEN, 74104) from HepG2 cells co-cultured for 20 hours with or without neutrophils, or with or without the neutrophil granule fraction (each in triplicate). RNA integrity number was determined using an Agilent Bioanalyzer 2100 (Agilent Technologies, Santa Clara, California, USA). On average, 40 million sequencing reads (100 bp paired end) of each sample were obtained using an Illumina 6000 Sequencer. The raw RNA-seq data has been deposited in the international NCBI SRA database as BioProject PRJNA876711 and PRJNA876578. Ingenuity Pathway Analysis: The sequencing data of HepG2 co-cultured with or without neutrophils were analyzed to get the candidates for functional blocking experiments. Quality control was performed by FastQC (Galaxy V.0.2.6+galaxy0) and Trimmomatic (Galaxy V.0.2.6+galaxy0). To remove Illumina adaptor sequences, the TruSeq3-paired-end file was used in Trimmomatic. The reads were mapped to the University of California Santa Cruz hg19 human genome by STAR (Galaxy V.2.6.0b-1). The raw read counts of genes were counted by feature Counts (Galaxy V.1.6.4+galaxy1) and the differential expression analysis was conducted by DESeq2 (Galaxy V.2.11.40.6). A list of gene transcripts satisfying FDR<0.05, fold change ±2, was input to IPA (Ingenuity Pathway Analysis, QIAGEN,). Integrins αM and 2 were selected as candidates from the top5 classical pathways obtained from IPA, from the list of genes defined as present in the plasma membrane in the gene ontology term. Bulk RNA-seq data analysis: To capture expression of genes commonly changed in HepG2 cells treated with neutrophil or granule fractions, both of sequencing data were analyzed. Low quality ends (less than phred score=30) and short read length (minimum length=30) was trimmed using PRINSEQ++ ^45^(version1.2). Trimmed reads were aligned to the hg38 genome reference using STAR^46^(v2.7.1), and subsequently RSEM (RNA-Seq by Expectation-Maximization) ^47^ was used to count reads mapping to the genes from Ensembl release 93. Normalized gene counts were evaluated using the R package DESeq2^48^. Heatmaps showed row-normalized relative gene expression z-scores across columns.

### Administration of protein to HepG2 cell homogenates

HepG2 cells were homogenized in 100 μL of PBS with a stick ultrasound device at 120 watts, 20 kHz output for 3 seconds for three times. Neutrophil myeloperoxidase (MPO), neutrophil elastase, recombinant proteins Serpin E2 (BioLegend, 769002) and Serpin A3 (R&D systems, 1295-PI) were administered for 5 minutes at indicated concentrations.

### Mouse model of alcoholic hepatitis

Alcoholic hepatitis was induced in male C57BL/6N mice aged 9-10 weeks (Charles River Laboratories) using the National Institute on Alcohol Abuse and Alcoholism (NIAAA) model of chronic plus binge ethanol feeding^32^. Lieber-DeCarli control and ethanol liquid diets were obtained from BioServ (Flemington, New Jersey, USA). After acclimatization to tube feeding with the control diet ad libitum for 5 days, mice received either the control liquid diet (paired-fed control group) or the ethanol liquid diet (ethanol-fed group) for 10 days. On day 11, mice were gavaged with 5 g ethanol/kg body weight and sacrificed 9 hours later. Liver tissue was harvested and frozen at −80°C. Then frozen liver tissue was homogenized and lysed with RIPA buffer (Thermo Fisher Scientific, 89900) containing a protease and phosphatase inhibitor cocktail (Thermo Fisher Scientific, 1861281). Extracted protein was assessed for ITPR2 by immune blotting. All animals received humane care in accordance with the Guide for the Care and Use of Laboratory Animals as adopted and promulgated by the National Institutes of Health. Animal studies were approved by Yale University’s Institutional Animal Care and Use Committee (IACUC).

### Statistical analysis

Data are expressed as mean ± SD or SEM of multiple independent experiments, unless stated otherwise. Statistical analyses were performed using the paired, unpaired Student’s t-test, or one-way analysis of variance with Tukey’s multiple comparison test where appropriate. Differences with p<0.05 were considered statistically significant. All statistical analyses were performed using GraphPad Prism 7 Software (GraphPad).

## Acknowledgements

We thank the Yale Liver Center for providing human blood from healthy controls. We also thank JIttima Weerachayaphorn for sharing her observations that neutrophils decrease ITPR2 in hepatocytes and that ITPR2 in liver is decreased in patients with Alcoholic Hepatitis, and we thank Mateus Guerra for assistance with conducting experiments and data analysis.Normal human hepatocytes were obtained from the Yale Liver Center or through the Liver Tissue Cell Distribution System, Pittsburgh, PA, which was funded by NIH Contract N01-DK-7-0004/HHSN267200700004C. This work was supported by grants from NIH (P01-DK57751, P30-DK34989, R01-DK114041, R01-DK112797, R01-AA028765), CAPES PrInt 88887.569334, CNPq 304451/2018-5, and FAPEMIG, and UFMG Liver center.

---

**Supplementary Figure 1.**
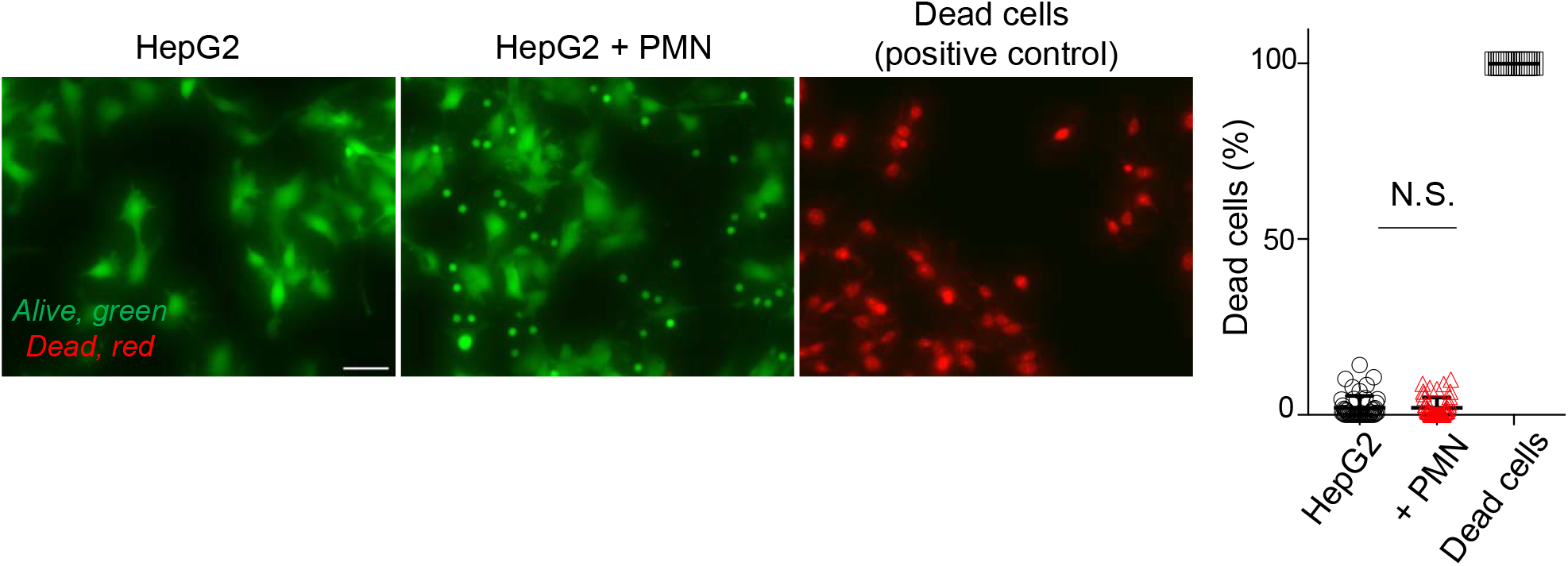
Human polymorphonuclear neutrophils (PMN) do not induce cell death in HepG2 cells, even after 40 hours of exposure. Live/dead cell analysis of HepG2 cells using double staining with calcein AM (green) and ethidium Homodimer-1 (EthD-1, red) with or without PMN for 40 hours. For positive control of dead cells, HepG2 cells were treated with 70% ethanol for 10 minutes. The representative image (*top*, scale bar: 50 μm) and quantification of dead cells (%) are shown (*right*). Data are mean ± SD, n=5-10 fields in 3 coverslips per each condition in 3 independent experiments. (N.S., no significant difference by unpaired t-test).

**Supplementary Figure 2.**
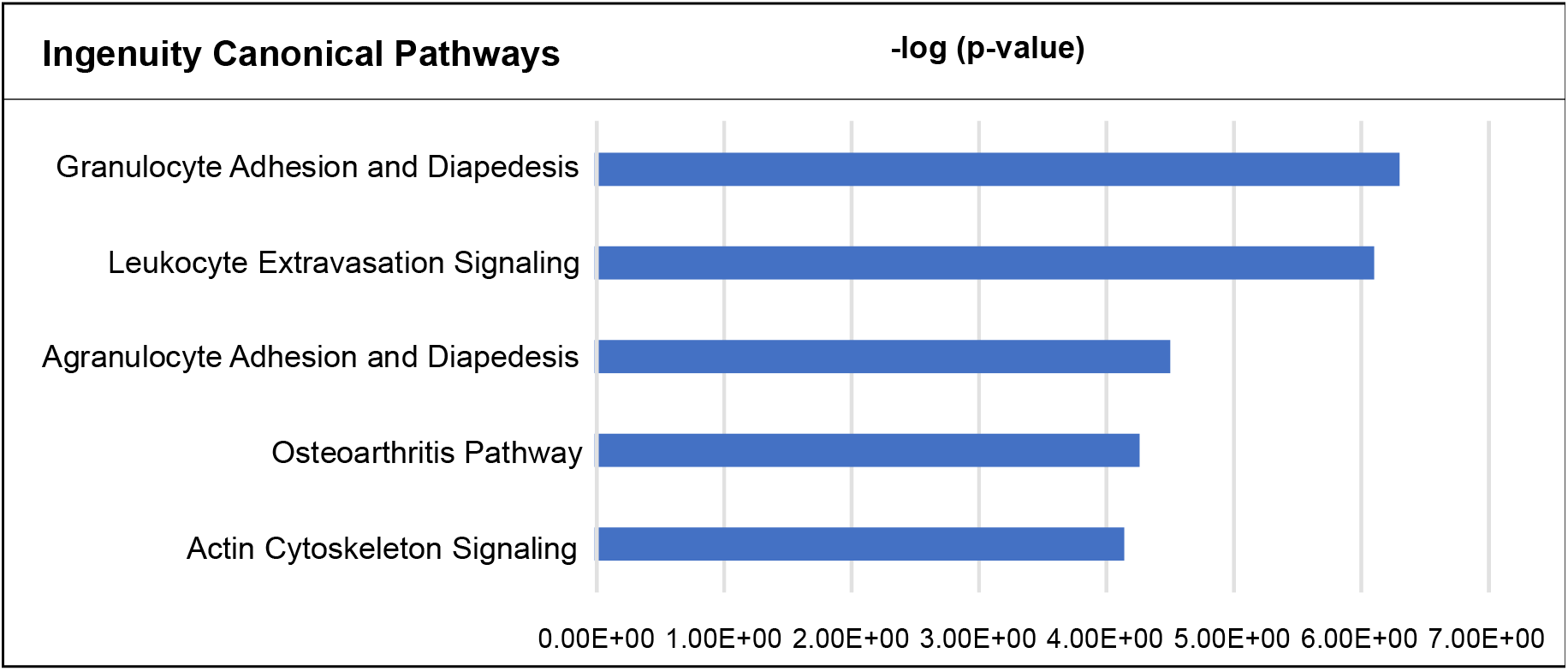
Top 5 canonical pathways derived from ingenuity pathway analysis (IPA) are shown. A list of gene transcripts satisfying false discovery rate (FDR)<0.05, fold change ±2, from RNA-seq data of HepG2 cells co-cultured with neutrophils for 20 hours, was analyzed. The values of -log(p-value) for each pathway are 6.294 (granulocyte adheision and diapedesis), 6.103 (leukocyte extravasation signaling), 4.488 (agranulocyte adhesion and diapedesis), 4.262 (osteoarthritis pathway), 4.146 (actin cytoskelton signaling) respectively.

**Supplementary Figure 3.**
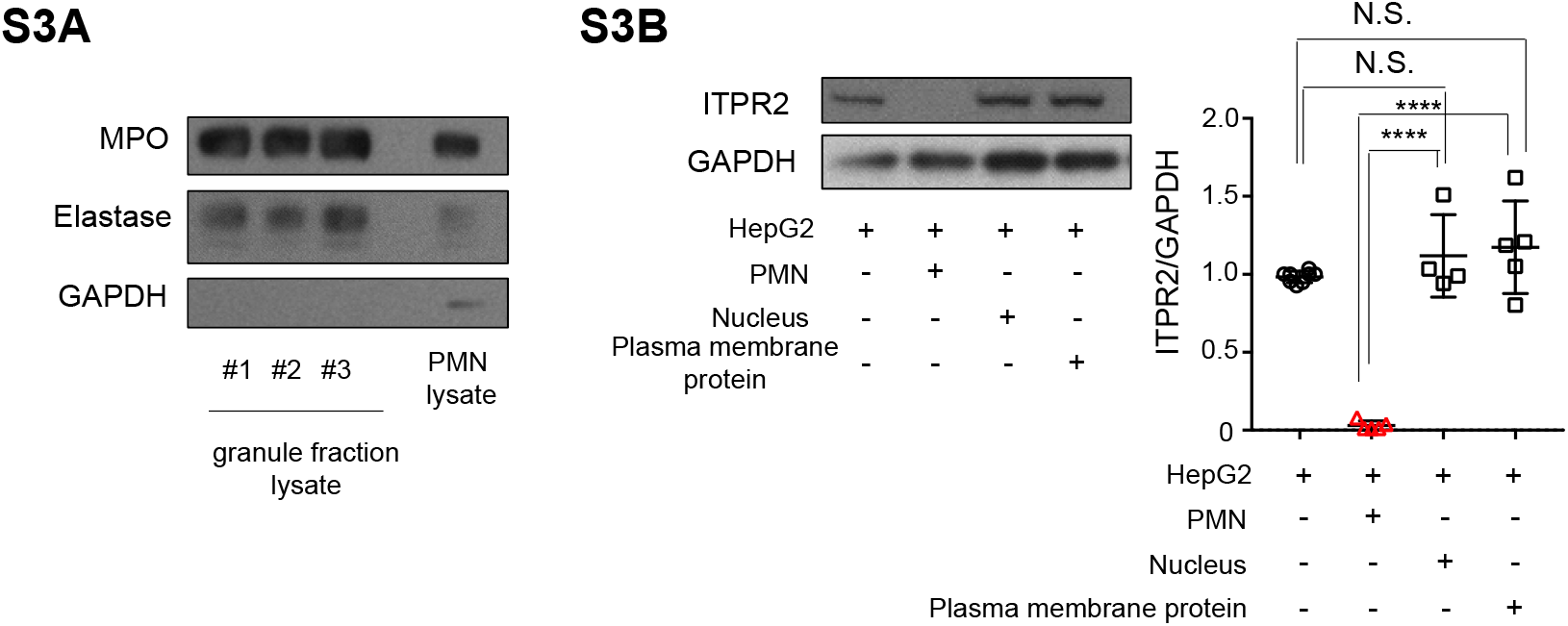
Granule of neutrophils contribute to the reduction of ITPR2 in HepG2 cells. **(a)** The granule fraction of human polymorphonuclear neutrophils (PMN) contains MPO and elastase. Immunoblots for myeloperoxidase (MPO) and elastase of granule fraction lysates extracted from PMN from three volunteers (#1-3) are shown. The rightmost lane shows the lysate of PMN as a positive control. **(b)** Neither the plasma membrane nor the nucleus of PMN decreases ITPR2 in HepG2 cells. Plasma membranes or nuclei were extracted from PMN using the Minute™ Plasma Membrane Protein Isolation and Cell Fractionation Kit. These fractions were administered to HepG2 cells for 20 hours and then the levels of ITPR2 were assessed by immunoblotting. Representative blots (*left*), and quantification (*right*) are shown. Data are mean ± SD, n=4-5. (N.S., no significant difference, ****p<0.0001 by one-way ANOVA).

**Supplementary Figure 4.**
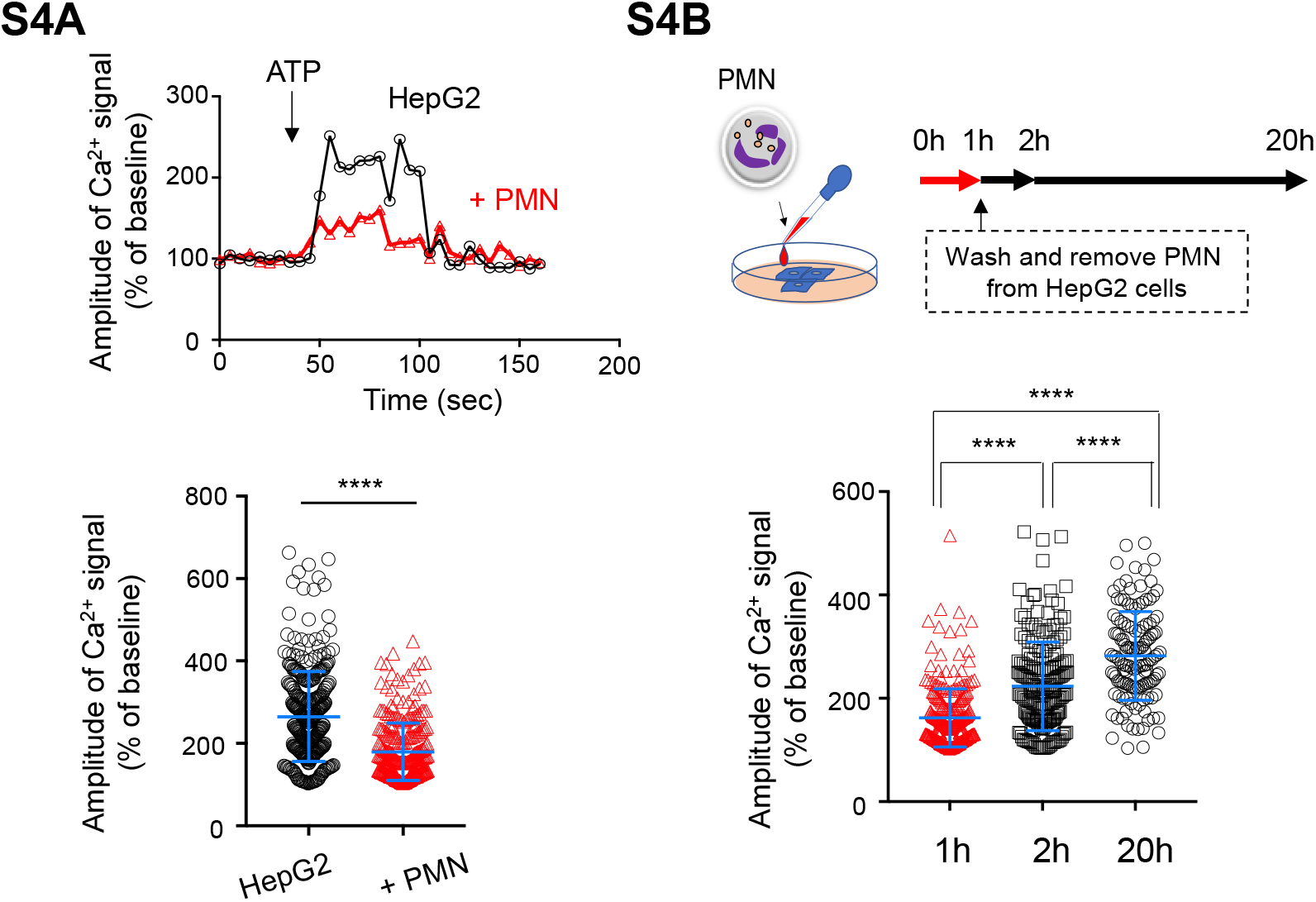
Calcium signaling is reversibly impaired in HepG2 cells exposed to neutrophils. **(a)** Human polymorphonuclear neutrophils (PMN) decrease the amplitude of calcium signals in HepG2 cells. Cells were pre-incubated for 1 hour with or without PMN, then loaded with 6 μM Fluo-4/AM for 30 minutes and stimulated with 20 μM adenosine triphosphate (ATP). Representative time course of amplitude of calcium signal (% of baseline) are shown measured in selected regions of interest of the cells using ImageJ. The right graph was results of quantification (6 coverslip of HepG2, n=330; 6 cover slips of HepG2 + PMN n=283, from 2 independent experiments, graphs are mean ± SD, ****p <0.0001, unpaired t-test). **(b)** After removal of PMN, calcium signals in HepG2 cells improve over time. HepG2 cells were loaded with 6 μM Fluo-4/AM for 30 minutes and stimulated with 20 μM ATP. The amplitude of calcium signals (% of baseline) were measured in selected regions of interest of the cells using ImageJ at 3 time points: when HepG2 cells were co-cultured with PMN for 1 hour, when PMN were completely washed and cultured for 1 or 19 hours (labeled as 2h and 20h). The graph shows the results of quantification (5-7 coverslip per each, cell number are n=266, 208, 135 from 2 independent experiments, graphs are mean ± SD, ****p<0.0001, one-way ANOVA).

**Supplementary Figure 5.**
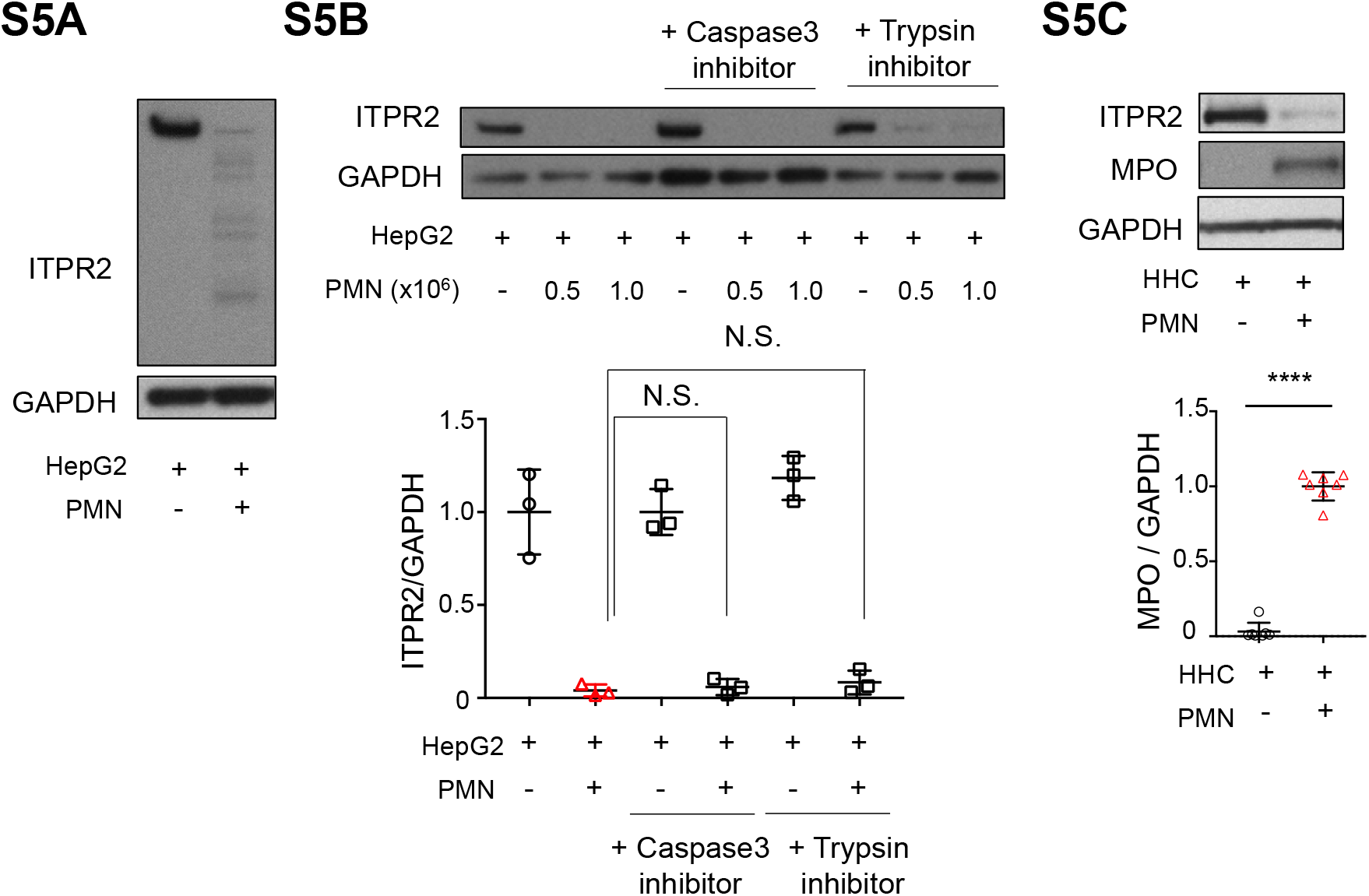
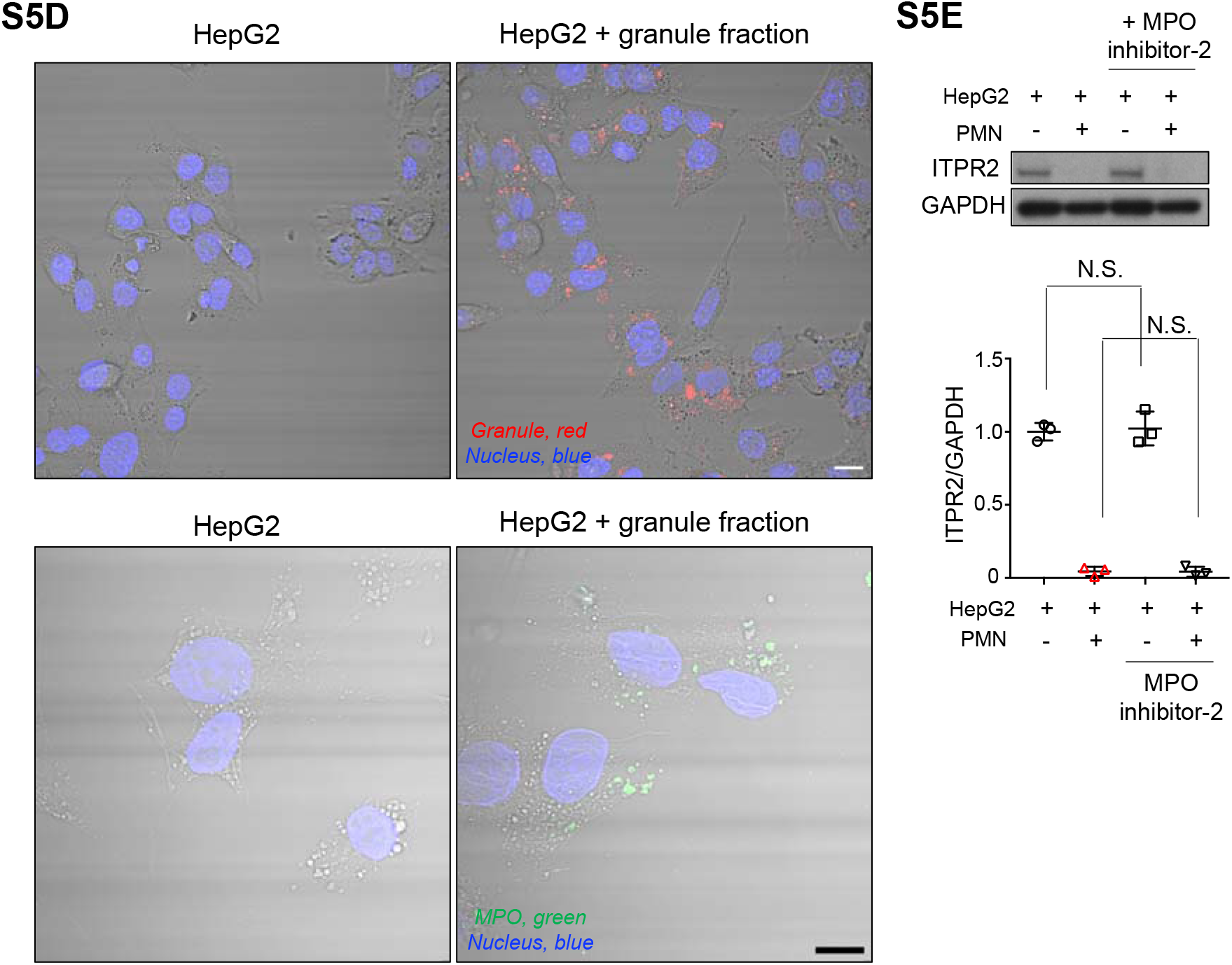
ITPR2 in HepG2 cells is degraded by neutrophils but not by caspase3, trypsin, or myeloperoxidase. **(a)** Longer exposure of the representative immunoblot in Fig.1B shows that the ITPR2 bands are laddered. **(b)** Human polymorphonuclear neutrophils (PMN)-induced loss of ITPR2 in HepG2 cells is not prevented by a trypsin inhibitor or a caspase 3 inhibitor. HepG2 cells were co-cultured with PMN for 20 hours with/without a caspase 3 inhibitor (50μM) or a trypsin inhibitor (AdaAhx3L3VS, 50nM). After co-culture with PMN, the ITPR2 levels in the HepG2 cells was assessed by immunoblotting. Representative blots of HepG2 cells co-cultured with 0.5 million and 1 million PMN are shown (*top*). Quantitative analysis (*bottom*) is shown only for HepG2 cells co-cultured with 1 million PMN. Data are mean ± SD, n=3 (N.S., no significant difference by one-way ANOVA). **(c)** Granule proteins are transferred from neutrophils into primary human hepatocytes (HHC). After co-culture with PMN for 20 hours, HHC were assessed for ITPR2 and myeloperoxidase (MPO) by immunoblotting. Representative blots (*top*) and quantification for MPO (*bottom*) is shown. Data are mean ± SD, n=7 from 3 independent experiments (****p<0.0001 by one-way ANOVA). **(d)** HepG2 cells take up components of the granule fraction from neutrophils. HepG2 cells were incubated with neutrophil-derived granule fraction stained with CellMask Orange fluorescent lipophilic membrane dye for 1 hour, washed, then observed by confocal microscopy. A representative images shows red granules incorporated in HepG2 cells (*top*, scale bar: 20 μm). In addition, HepG2 cells were incubated with non-stained granule fraction for 1 hour and observed by confocal microscopy after immunostaining for anti-MPO antibody (*green*). A representative images of HepG2 cells are shown (*bottom*, scale bar, 20 μm). **(e)** MPO inhibitor-2 (5-Fluoro-1H-indol-3-yl butanamide) does not prevent the decrease in ITPR2 in HepG2 cells co-cultured with PMN. After co-culturing with PMN with or without 5 μM MPO inhibitor-2 for 1 hour, the ITPR2 levels in HepG2 cells were assessed by immunoblotting. Representative blots (*top*) and quantification (*bottom*) are shown. Data are mean ± SD, n=3 (N.S., no significant difference by one-way ANOVA).

**Supplementary Figure 6.**
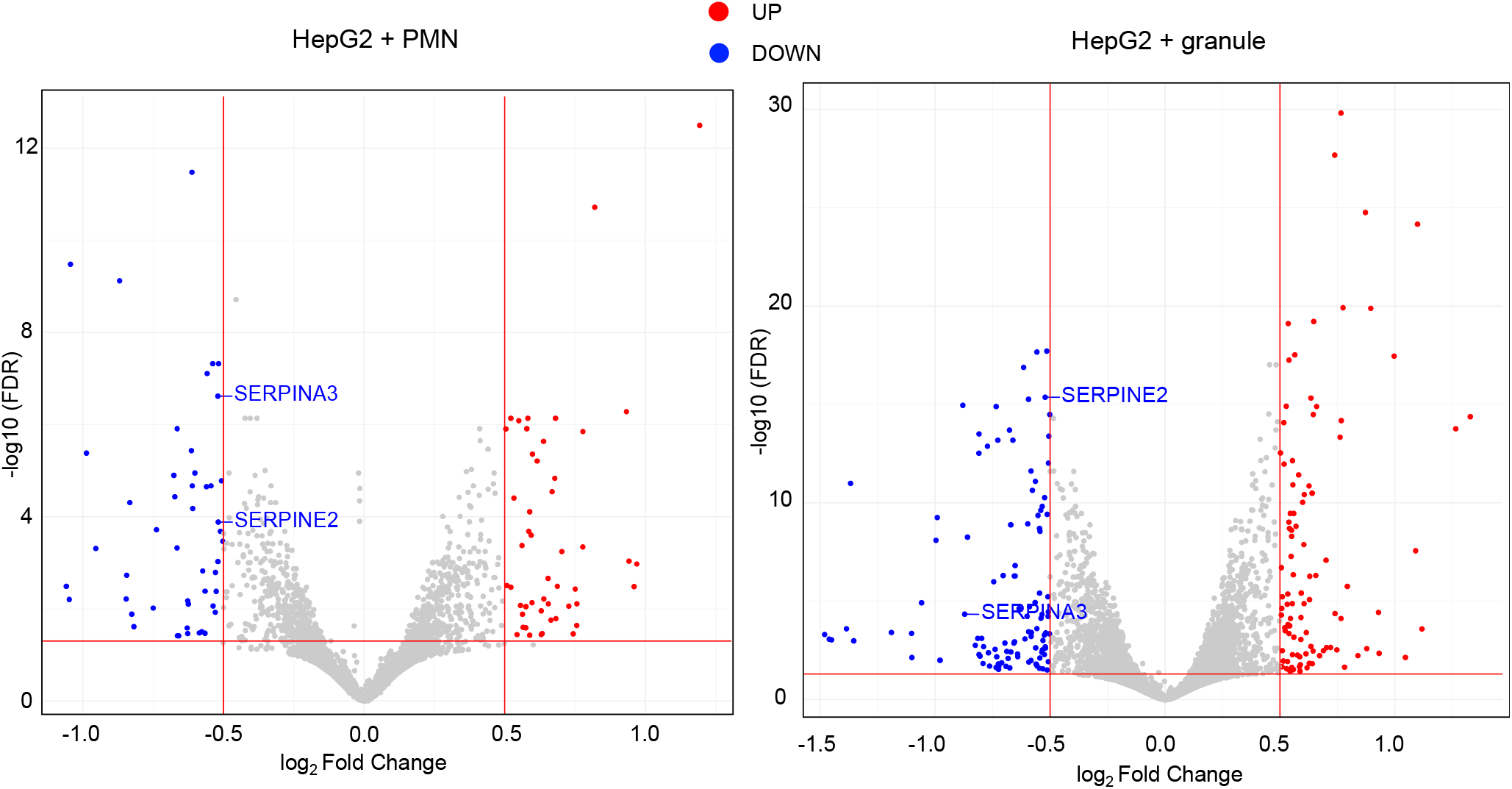
Volcano plots of up- or down-regulated differentially expressed genes from RNA-seq for HepG2 cells treated with neutrophils (HepG2 + PMN, *left*) or extracted granule fraction (HepG2 + granules, *right*) for 20 hours, compared to control respectively (triplicate per each). Grey dots represent genes which are not differentially expressed, red dots (UP) represent the upregulated genes, and the blue dots (DOWN) represent the downregulated genes. The threshold for the analysis was set at false discovery rate (FDR)<0.05 and log Fold change < −0.5 or >0.5 (vs control). Serpin E2 and Serpin A3 are included as significantly decreased genes in both analyses.

**Supplementary Figure 7.**
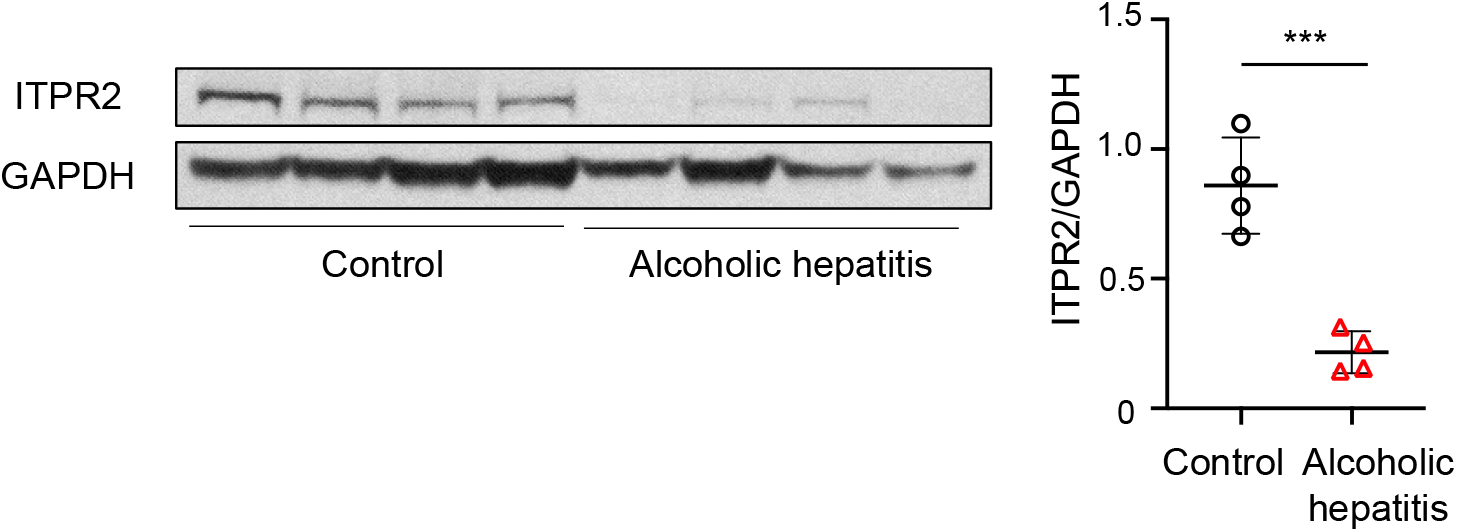
ITPR2 is decreased in the liver in a mouse model of alcoholic hepatitis. The levels of ITPR2 were assessed by immunoblotting in hepatic lysates from control mice and from mice model for alcoholic hepatitis. Representative blots (*left*) and quantification (*right*) are shown. Data are mean ± SD, n=4 per each, (***p<0.001 by unpaired t-test).

**Supplementary table1.**
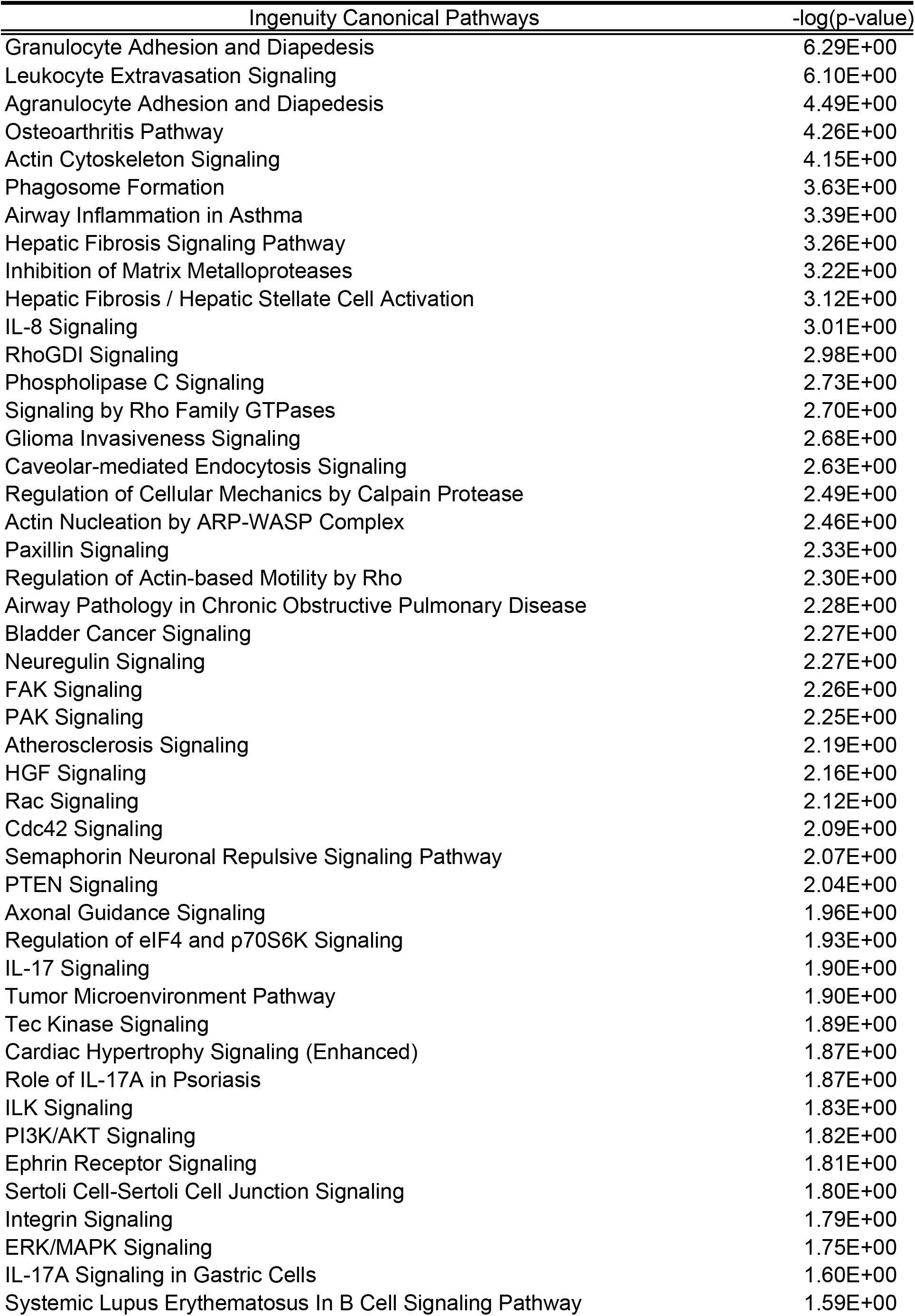
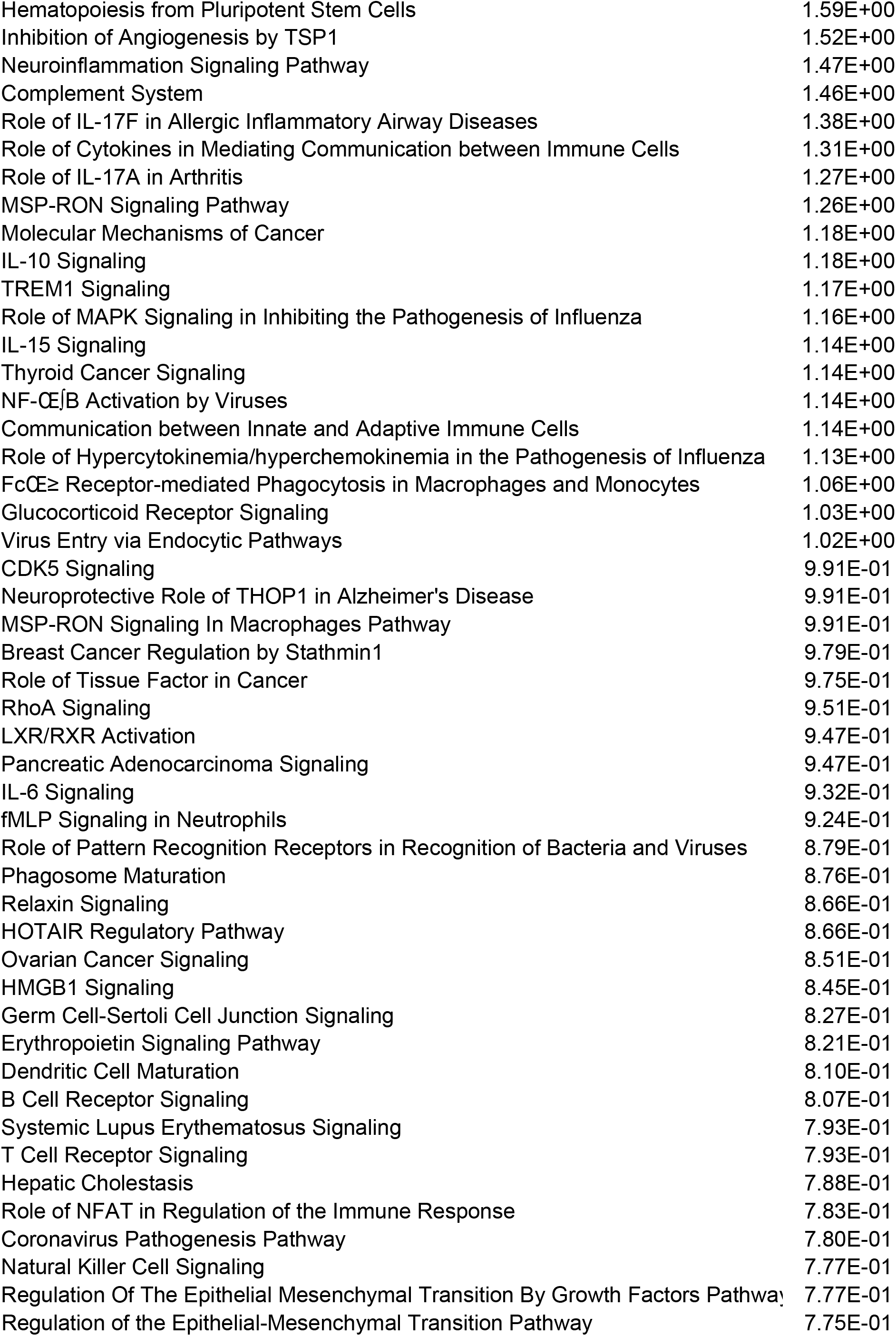
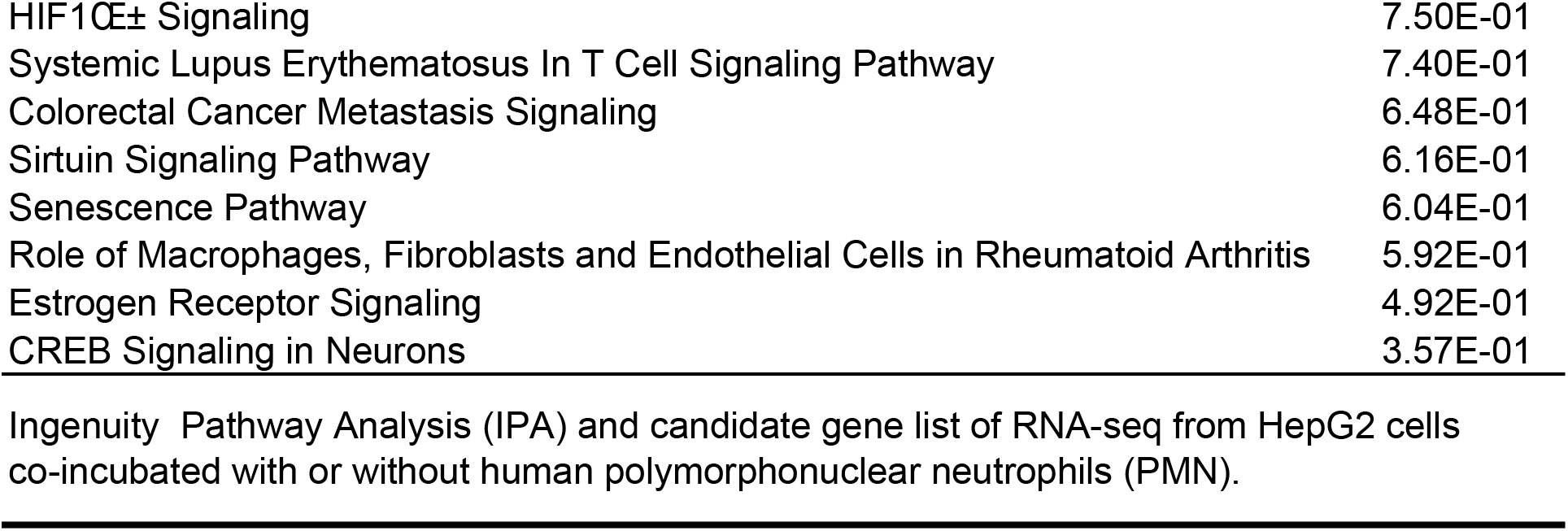

